# Direct visualization of native CRISPR target search in live bacteria reveals Cascade DNA surveillance mechanism

**DOI:** 10.1101/589119

**Authors:** Jochem N.A. Vink, Koen J.A. Martens, Marnix Vlot, Rebecca E. McKenzie, Cristóbal Almendros, Boris Estrada Bonilla, Daan J.W. Brocken, Johannes Hohlbein, Stan J.J. Brouns

**Author notes:** Corresponding authors: Brouns, S.J.J., Hohlbein, J.

## Abstract

CRISPR-Cas systems encode RNA-guided surveillance complexes to find and cleave invading DNA elements. While it is thought that invaders are neutralized minutes after cell entry, the mechanism and kinetics of target search and its impact on CRISPR protection levels have remained unknown. Here we visualized individual Cascade complexes in a native type I CRISPR-Cas system. We uncovered an exponential relationship between Cascade copy number and CRISPR interference levels, pointing to a time-driven arms race between invader replication and target search, in which 20 Cascade complexes provide 50% protection. Driven by PAM-interacting subunit Cas8e, Cascade spends half its search time rapidly probing DNA (∼30 ms) in the nucleoid. We further demonstrate that target DNA transcription and CRISPR arrays affect the integrity of Cascade and impact CRISPR interference. Our work establishes the mechanism of cellular DNA surveillance by Cascade that allows the timely detection of invading DNA in a crowded, DNA-packed environment.

**One sentence summary:** The results from *in vivo* tracking of single CRISPR RNA-surveillance complexes in the native host cell explain their ability to rapidly recognize invader sequences.

## Introduction

RNA-guided CRISPR-Cas surveillance complexes have evolved to specifically and rapidly recognize sequences of previously catalogued mobile genetic elements (MGEs) (Marraffini, 2015). Target DNA recognition depends on CRISPR RNA (crRNA) – DNA complementarity and on the presence of a protospacer adjacent motif (PAM), a short nucleotide sequence flanking the target site (Deveau et al., 2008; Mojica et al., 2009). To work effectively, the complexes need to find their targets fast enough to prevent an MGE from becoming established in the cell, which can occur within minutes upon cell entry (Shao et al., 2015). Target search inside a cell faces a multitude of challenges: Firstly, cells are packed with DNA, and crRNA surveillance complexes need to find the needle in a haystack before an invading element takes control of the cell. PAM scanning and crRNA-seed interactions with the target have been suggested to speed up the search process by drastically reducing the number of potential target sites in the genome (Gleditzsch et al., 2018; Jones et al., 2017). Several studies have shown that crRNA-effector complexes spend more time probing PAM rich regions, which is indicative of its function as the first recognition site (Globyte et al., 2018; Redding et al., 2015; Sternberg et al., 2014). In the *Escherichia coli* Type I-E CRISPR-Cas system, the crRNA-effector complex Cascade recognizes six PAMs with high affinity (Leenay et al., 2016) suggesting that Cascade scans hundreds of thousands of PAM motifs in the host genome, which is only effective when this interaction is sufficiently fast. A second challenge is posed by the action of other proteins present in the cell such as DNA binding proteins, DNA or RNA polymerases that may interfere with target search and formation of target bound crRNA complexes (Jones et al., 2017; Vigouroux et al., 2018). Some invading MGEs even use specialized anti-CRISPR proteins to inhibit crRNA-effector complexes and impair the target search process (Bondy-Denomy et al., 2015; Pawluk et al., 2014). A third challenge that microbes face is to produce appropriate levels of Cascade complexes loaded with one particular crRNA to provide protection against a single invading element. While adding more and more spacers to CRISPR arrays will have the benefit of recognizing many invaders, the tradeoff is that long CRISPR arrays will dilute the number of Cascade complexes loaded with a particular crRNA, potentially decreasing the CRISPR response against that target. These cellular challenges raise the question how Cascade can navigate the crowded cell sufficiently fast to find DNA targets, and how many copies of Cascade are required to do so.

Here, we report the visualization of single-molecule Type I-E Cascade complexes in a native *E. coli* CRISPR-Cas system *in vivo*. We found that the probability of successful CRISPR protection depends exponentially on Cascade copy numbers, which leads to a time-driven arms race model between Cascade target search and invader replication. The localization of Cascade shows the complex is enriched inside the nucleoid. We determined that 60% of the Cas8e subunit is incorporated into Cascade complexes and that Cascade DNA probing is very rapid (∼ 30 ms) and is driven by Cas8e. Furthermore, transcription of targets and CRISPR arrays reduce the amount of functional complexes in the cell. Our work sheds light on target search and dynamical assembly of Cascade complexes in their native cellular environment, and describes how these processes impact CRISPR protection levels.

## Results

### Visualizing Cascade abundance and target search at the single-molecule level

To investigate how microbes deal with these challenges at the cellular level we used a intracellular single-particle tracking Photo-Activated Localization Microscopy (sptPALM) (English et al., 2011; Manley et al., 2008), a technique capable of following the movement and abundance of individual fluorescently-tagged proteins in cells with high precision. By genetically fusing a photoactivatable fluorescent protein (PAmCherry2, (Subach et al., 2009)) to the N-terminus of Cascade-subunit Cas8e (Figure 1A), which was the only subunit for which labeling had no influence on the CRISPR interference ability of this strain (Figure 1B), we were able to monitor the mobility and abundance of Cascade complexes in *E. coli* cells.

**Figure 1:**
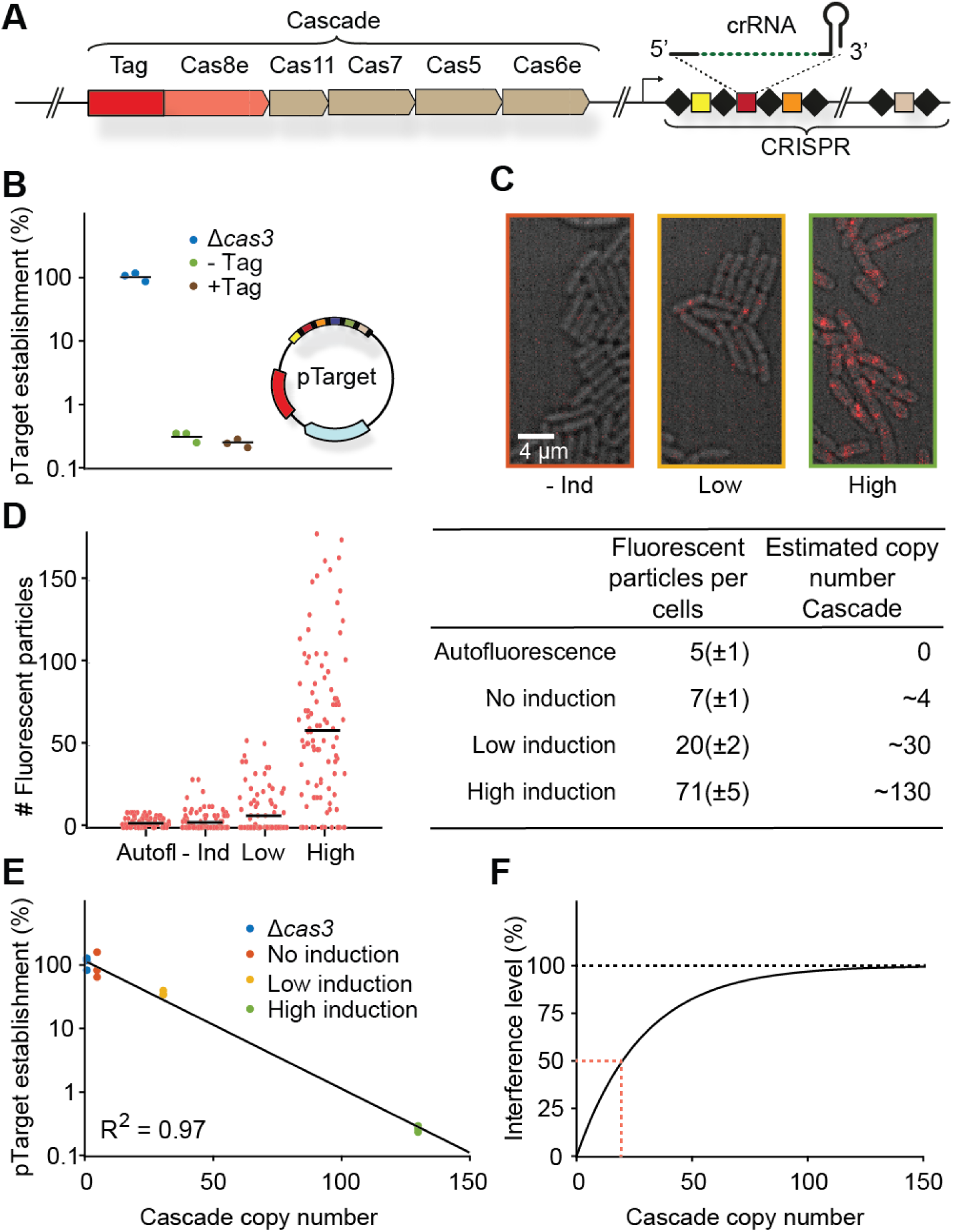
Cascade copy number vs CRISPR protection. (A) Chromosomal locus of the Cascade subunits and integration site of the photoactivatable fluorescent protein upstream of *cas8e.* (B) pTarget establishment, calculated from the ratio of transformation of pTarget/pGFPuv, is a measure for the interference level of the CRISPR system. To test whether tagged Cascade complexes were able to function normally, we compared the tagged strain to the untagged and the *Δcas3* strain. pTarget (bottom right) contains protospacers for all spacers in the K12 genome (colored, not all depicted) and are flanked by a 5’-CTT-3’ PAM (black bars). (C) Overlay of brightfield image of cells (grey) and single molecule signal (red) from a single representative frame for different induction levels. (D) Number of fluorescent particles measured in each cell plotted for different levels of Cascade expression (left). The mean number of fluorescent particles (± standard deviation; table left column) was converted to a Cascade copy number (table right column, Methods). (E) pTarget establishment plotted for different copy numbers of Cascade. The data points were fitted with an exponential decay function. *pTarget establishmentnt* = *e*^−*an*^, where *n* equals Cascade copy number and *a* the fitted coefficient. In our model 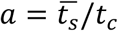. (F) The fitted exponential decay fitted on the left converted into an interference level (*Interference level* = 1 − *pTarget establishmentnt*). Indicated in red (dashed) is the amount of Cascade copies required for 50% interference.

### Twenty Cascade complexes provide 50% CRISPR protection

We first wanted to link the copy number of Cascade to successful target search, and established an assay that measures the level of CRISPR protection in cells at the time of cell entry by a mobile genetic element (MGE). In this assay all Cascade complexes present in the cell must be able to target the incoming MGE and Cascade target search has to be rate limiting. To meet the first requirement, we constructed a high copy plasmid (pTarget; Figure 1B) containing target sites for all 18 spacers found in the genomic arrays of *E. coli* K12, such that all Cascade complexes would be targeting the incoming plasmid. Secondly, we ensured that Cascade copy numbers were rate limiting (Majsec et al., 2016) by equipping cells with a low copy plasmid expressing the nuclease Cas3 (pCas3, adapted from (Westra et al., 2010)).

We achieved different expression levels of Cascade expression in the cell by tuning the expression of the native regulator LeuO (Westra et al., 2010) (Figure 1C). The copy numbers of Cascade were estimated from the number of fluorescent particles present in the cell under these varying levels of LeuO induction, taking complex assembly (see following section), growth rate (Table S1) and maturation time of PAmCherry into account (Figure 1D; Methods). We found that the average number of Cascade complexes per cell in the absence of LeuO induction was low (∼4 copies) and that copy numbers increased more than 30-fold for the highest induction level (∼130 copies). We measured the interference ability under these conditions by determining the probability that pTarget becomes established in a cell. We observed that establishment of pTarget decreases sharply with increasing copy numbers of Cascade (Figure 1E). However, even with 130 Cascade complexes present, we still observed a level of pTarget survival (∼0.5%).

To explain these observations, we modelled the probability that an invading MGE becomes established in the cell depending on the number of Cascade complexes that target this specific MGE. The model is based on multi-copy plasmids and phage systems, where the DNA clearance is most likely to occur at when an invader enters as a single copy, as the concentration of invading DNA increases over time. Therefore, depending on the invader and the level of CRISPR interference, there will be a critical time point (*t*_c_) beyond which the invader is permanently established inside the cell and can no longer be cleared (Severinov et al., 2016). Our model describes the probability that it takes a certain copy number of proteins (*n*) each with an average search time 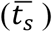 to find the target before *t*_c_ is reached. Our model accurately predicted that pTarget establishment decreases exponentially with increasing copy numbers of Cascade (Figure 1E, Methods).

When we translated these establishment probabilities in interference levels, we could deduce that around 20 Cascade complexes are required to reach a CRISPR interference level of 50% (Figure 1F). It becomes very unlikely for the CRISPR system to destroy multiple genetic copies of the MGE if it has failed to destroy the single copy that was present at the start before replication. Therefore, we can approximate *t*_c_, with the replication time of the plasmid in the absence of copy number control (∼3 min, (Olsson et al., 2003a)), which allows us to retrieve an estimated search time of ∼90 minutes for one Cascade complex to find a single target in the cell (Methods).

To summarize, we found a direct relation between the number of Cascade complexes and the establishment probability of an MGE. The native *E. coli* system requires 20 Cascade complexes loaded with a cognate crRNA to obtain 50% CRISPR interference levels. This relation depends on the replication rate of the invading MGE and the average search time of a single complex and demonstrates the importance of rapid target search on CRISPR interference ability.

### The majority of Cas8e assembles into the Cascade complex

To quantify the dynamics of target search, we traced the diffusion paths of thousands of individual complexes in the bacterial cell (Figure 2A; Supplementary Video). The apparent diffusion coefficient *D*\*, a measure for mobility, of Cascade was calculated by extracting the displacement of each fluorescent particle for four consecutive 10 ms steps, allowing us to investigate the abundance, mobility and behavior of individual complexes and subunits in the cell. To minimize the influence of spurious autofluorescent particles in *E. coli* (Floc’h et al., 2018), we used expression levels with the highest estimated Cascade copy numbers (∼130 copies, high induction; Figure 1D).

**Figure 2:**
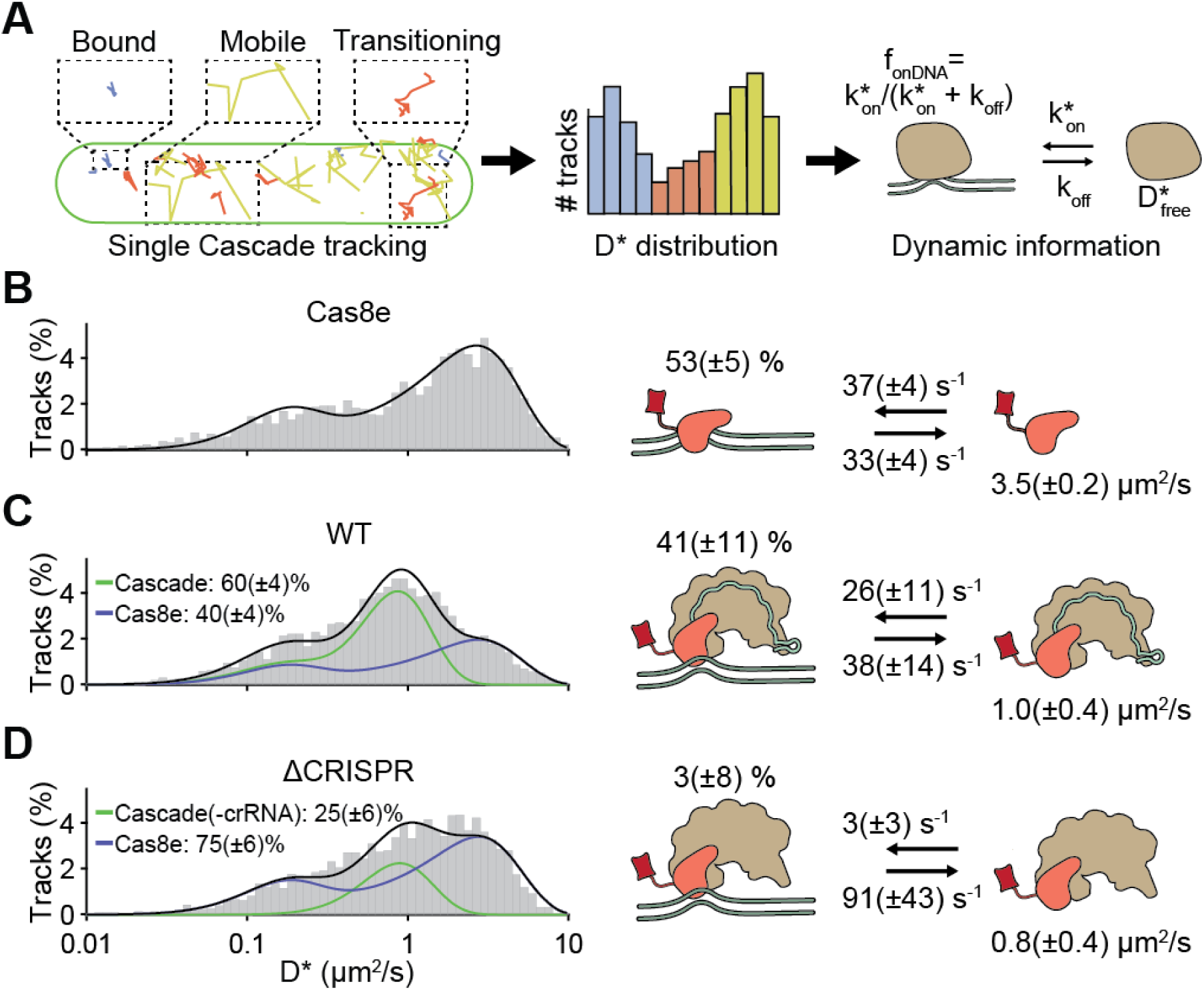
Diffusion behavior of Cas8e and Cascade. **(A)** Tracks recovered from a single cell of the WT strain (left). Proteins can be bound to DNA (blue), be freely diffusing (yellow) or transitioning between bound and mobile states (orange) within a track. The analytical diffusion distribution analysis (DDA; right) extracts kinetic information (pseudo-first order on-rate 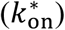, off-rate (*k*_off_) and the apparent free diffusion coefficient (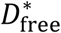)) from *D\** distributions, which further allows the calculation of fraction DNA bound (*f*_onDNA_). **(C-E)** *D\** distributions for **(B)** Cas8e, **(C)** Cascade and **(D)** ΔCRISPR strain. Total (black), Cas8e (blue) and Cascade (green) fractions fits are indicated by lines. Parameters for Cas8e (B) and Cascade (C-D) are depicted on the right. Parameters from (B) were used to fit the Cas8e fraction in (C) and (D). Error estimation is based on bootstrapping (± standard deviation). See also Figure S1 and S2.

To distinguish diffusion of Cascade complexes from monomeric Cas8e subunits, we first measured the diffusion of the tagged Cas8e fusion protein in a strain lacking genes of the other four Cascade subunits in the genome (Cas11, Cas7, Cas5, and Cas6e). Based on the role of Cas8e in non-specific DNA binding (Brown et al., 2018; Jore et al., 2011; Sashital et al., 2012), we expected to find mobile and DNA-bound populations of Cas8e. However, we were unable to describe the data accurately by static two-state models of non-interconverting fractions (Figure S1). We therefore hypothesized that rapid DNA binding and unbinding events of Cascade on a timescale similar to the framerate (∼10-40 ms) would lead to time-averaging of a mobile state (high *D\** values) and a DNA-bound state (low *D\** values), giving rise to intermediate *D*\* values (Figure 2A and S2). We accounted for these events by developing a generally applicable analysis method called analytical Diffusion Distribution Analysis (analytical DDA), which is useful for proteins with fast transitioning kinetics between states with different diffusion coefficients, such as DNA-interacting proteins. This method allows us to extract quantitative information on DNA binding kinetics (Figure S2), and enables the study of fast transition rates previously inaccessible to sptPALM (Methods).

When we applied the analytical DDA on the Cas8e diffusional data, we retrieved an average residence time of ∼30 ms on DNA and a similar average time spent (∼30 ms) rapidly diffusing (D* ∼3.5 μm^2^/s, as expected for a protein of 82 kDa; Methods), indicating that Cas8e is bound to DNA for ∼50% of the time (Figure 2B). The *D\** distribution of Cas8e then allowed us to extract the diffusion behavior of the Cascade complex as a whole. We estimated the fraction of free Cas8e and Cascade-containing Cas8e at 40% and 60%, respectively (Figure 1D). This finding suggests that Cas8e is produced in excess (Westra et al., 2010) or somehow involved in a dynamic interaction with the core Cascade subunits (crRNA, Cas11, Cas7, Cas5, Cas6e) (Jore et al., 2011; Sashital et al., 2012).

Surprisingly, we found that the DNA binding kinetics of Cascade were similar to Cas8e alone, indicating that Cas8e is an important driver of DNA probing characteristics of the Cascade complex. Furthermore, the DNA probing events take on average ∼30 ms and are thereby considerably faster than the 0.1-10 s that have been reported for *in vitro* studies previously (Brown et al., 2018; Redding et al., 2015; Xue et al., 2017). As expected, we found a smaller diffusion coefficient for unbound Cascade complexes (∼1.0 μm^2^/s) (Methods) due to their larger size. Together, our analysis shows that more than half of the Cas8e protein population is part of intact Cascade complexes, and that the DNA interacting behavior of Cascade is largely determined by the properties of Cas8e.

To investigate the role of crRNAs in Cascade complex assembly, we deleted all CRISPR arrays in the K12 genome (ΔCRISPR). The resulting diffusion behavior can be described by fractions of free Cas8e and with Cascade-like diffusion behavior (Figure 2D) that almost entirely lacks interaction with DNA (*f*_onDNA_ = 3%). This indicates that although Cascade (sub)complex formation does not strictly require the presence of crRNA (Beloglazova et al., 2015; Brouns et al., 2008), Cascade assembly is greatly enhanced by crRNA. Taken together, the majority of Cas8e proteins are incorporated in Cascade complexes in the presence of crRNA, and this gives Cascade DNA interacting properties.

### Cascade is enriched but not exclusively present in the nucleoid

Not all potential DNA interaction sites in the host chromosome might be accessible to Cascade. The host DNA is concentrated in the middle of the cell in the nucleoid and is very compact which excludes large complexes such as ribosomes (Mondal et al., 2011). Nucleoid exclusion would reduce the amount of DNA available for scanning and increase the amount of freely diffusing Cascade complexes. To investigate whether the DNA-bound fraction is governed by affinity properties of Cascade for DNA rather than a restricted search space outside the DNA-containing nucleoid region, we studied the spatial distribution of Cascade localizations. Nucleoid-excluded ribosomes are enriched away from the central long axis of the cell (Sanamrad et al., 2014). For Cascade, we found a homogeneous spatial distribution throughout the cell (Figure 3A), indicating that Cascade is small enough to freely scan the nucleoid for target sites.

**Figure 3:**
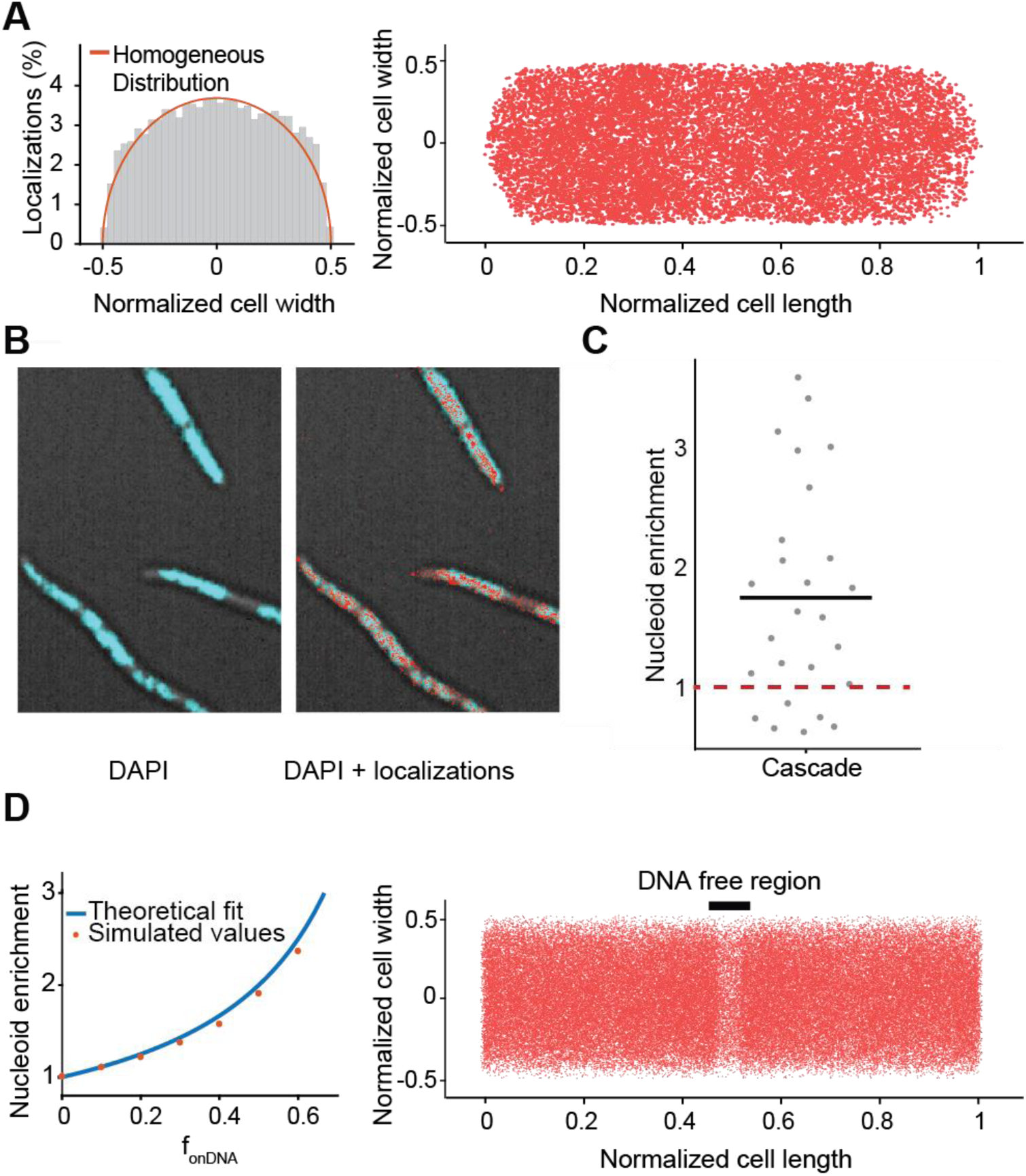
Cascade localization inside the cell. **(A)** Localization of Cascade in the cell. Left: Distribution of Cascade over the cell width of the cell from Cascade (n = 33 cells; 15428 localizations): in orange is indicated the expected distribution in case of a homogeneous localization within the cell. Right: same localizations plotted within dimensions of single cell in which the cell length and cell width of each cell was normalized. **(B)** Overlay of DAPI fluorescence and brightfield image (left) with Cascade localizations (right) in cephalexin treated cells. **(C)** The nucleoid enrichment in WT strain (27 subregions in 18 cells). The average ratio is indicated with a black bar. The expected ratio if Cascade has no interaction with DNA is indicated in red (dashed). **(D)** Relation between DNA bound fraction and nucleoid enrichment. Left: A theoretical relation between nucleoid enrichment and DNA bound fraction was derived (Methods) and compared to simulated values for different amounts of f_onDNA_. Right: Localizations of simulated Cascade proteins (n =50.000) diffusing through part of an elongated cell are plotted on top of long cell axis. A DNA-free region (black bar) is visible due to enrichment of Cascade binding to DNA in nucleoid regions. Simulations of particles were performed with off-rate of 38 s^-1^ and an on-rate of 26 s^-1^ to reach a nucleoid enrichment of 1.8, similar to the average that was found for Cascade.

We furthermore used the spatial distribution of Cascade to extract quantitative information on the DNA-bound fraction. To that purpose, we created a DNA-free environment in the cell by adding cephalexin (Reyes-Lamothe et al., 2014). This antibiotic affects cell wall synthesis and causes cells to elongate, forming DNA-free cytoplasmic space between nucleoids without condensing the nucleoid (Figure 3B). The time Cascade is bound to DNA is inherently linked to the relative amount it spends in DNA-free and DNA containing regions. Therefore, by calculating the relative amount of localizations in both regions (Enrichment Factor; *EF*) we can extract the fraction of time spent on DNA independently from the DDA analysis. Cascade was only moderately enriched (*EF* of 1.8 ± 0.2 fold) in the nucleoid regions (Figure 3C), indicating that Cascade spends a considerable amount of time diffusing in the cytoplasm while not associated with DNA. From the enrichment factor, the fraction of Cascade complexes bound to DNA can be approximated to 45% (Figure 3D; for derivation see Methods). This value is consistent with the ∼ 50% value we extracted from the DDA distribution of Cascade (Figure 2C). However, it strongly contrasts other DNA binding proteins such as Fis and RNA polymerase, which show a much higher nucleoid enrichment (Reyes-Lamothe et al., 2014; Stracy et al., 2015). The above findings indicate that Cascade inherently spends more time freely diffusing the cell and that this is caused by the nature of DNA-Cascade interactions and not by size-based nucleoid exclusion, as is the case for ribosomes (Sanamrad et al., 2014). Therefore, we decided to study the nature of the DNA interactions in more depth.

### Cascade-DNA interactions are not only PAM-dependent

Next, we assessed how PAM interactions contributed to DNA binding by introducing mutation G160A in the Cas8e subunit which abolishes the interaction with the PAM (Hayes et al., 2016). This G160A mutation decreased the fraction of DNA-bound Cascade from 41 ± 11 to 28 ± 6% (Figure 4A) without fully inhibiting DNA binding, suggesting that PAM-independent interactions (Van Erp et al., 2015; Hayes et al., 2016; Xiao et al., 2017) play a role in DNA probing as well. To assess the contribution of these different types of interactions to the average DNA residence time found previously, we measured the persistence of Cascade-DNA interactions by increasing the dark time between exposures (Figure 4B). Our data showed that sustained binding events at longer time scales (100 – 250 ms) were more frequently observed for WT Cascade than for the PAM binding mutant complex Cascade-Cas8e_G160A_ (Figure 4C). Together with the increased off-rate of the mutated complex (Figure 4A), this finding demonstrates that PAM-dependent interactions of Cascade with DNA last longer than PAM-independent interactions.

**Figure 4:**
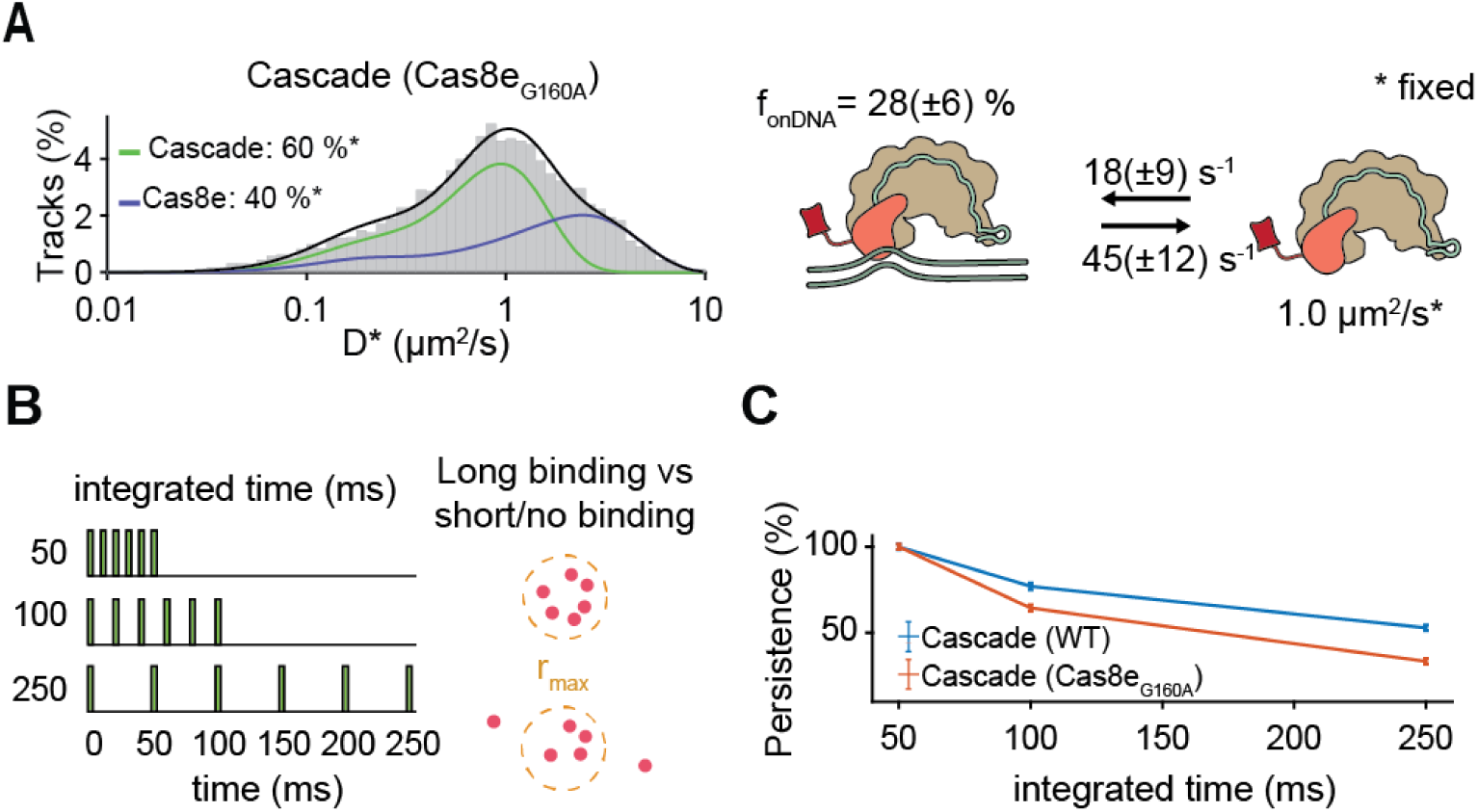
PAM-dependent and PAM-independent DNA probing. **(A)** *D\** distributions for Cascade and Cas8e with a mutation (G160A) deficient in PAM binding. To compare kinetic rates, we assumed that the relative Cas8e-Cascade fractions and the diffusion of free Cascade and Cas8e were not altered by the mutation and those values were fixed. **(B)** The relative amount of long binding events (6 consecutive localizations within r_max_: 1 pixel (0.128 μm) of the mean position) for WT and PAM binding mutant Cascade normalized to 50 ms integration time. Error estimation in (A) and (C) is based on bootstrapping (± standard deviation).

### Target DNA binding is influenced by the cellular environment

After establishing intrinsic DNA probing characteristics of Cascade, we next investigated its diffusion behavior in the presence of targets (Figure 5). To prevent target DNA degradation by Cas3 nucleases, we deleted the *cas3* gene and verified that the deletion did not alter Cascade diffusion behavior (Figure S3). To verify that all Cascade complexes could bind a target, we measured the copy number of pTarget to be ∼ 400 copies/cell (Figure S4). As the native *E. coli* CRISPR arrays contain 18 spacers, this resulted in ∼7000 target sites per cell which far outnumbers Cascade copy numbers under our growth conditions (∼130, Figure 1D).

**Figure 5:**
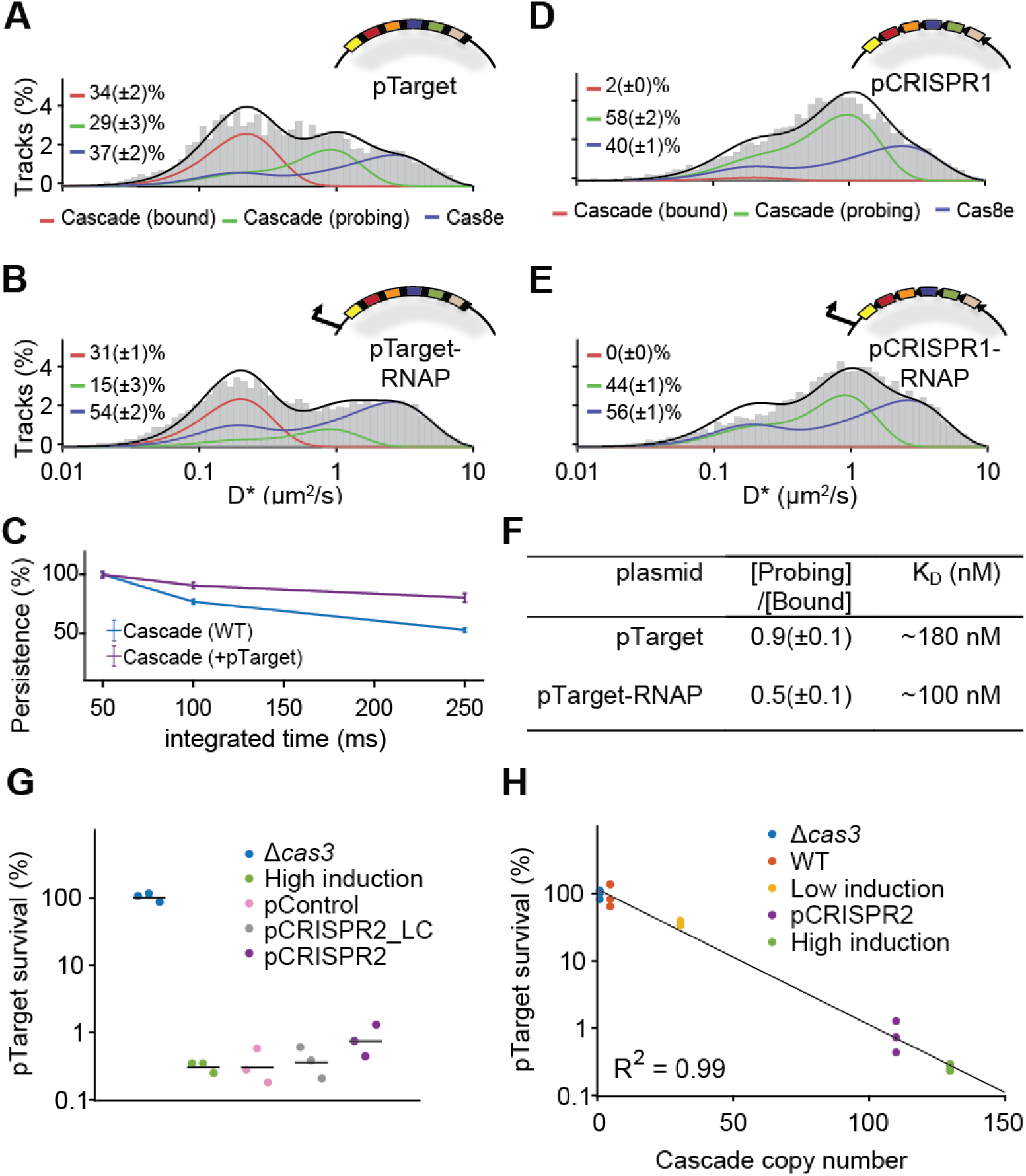
Cascade - DNA interactions in the presence of targets. (A and B), *D*\* distribution for the Δ*cas3* strain carrying pTarget (A) and pTarget-RNAP (B). pTarget contains protospacers for all spacers in the K12 genome (colored, not all depicted) and are flanked by a 5’-CTT-3’ PAM (black bars). Cascade (probing) (green) and Cas8e (blue) fractions were fitted with parameters from Figure 1C and 1D, and a new target-bound fraction (Cascade (bound)) was introduced as a single diffusion state (*D*\* = 0.06 μm^2^/s (+σ^2^/t); red). (C) The abundance of sustained binding events as in Figure 3C, but for WT and pTarget-carrying cells. (D and E), *D*\* distribution for the Δ*cas3* strain carrying pCRISPR1 (D) and pCRISPR1-RNAP (E). pCRISPR1 contains the same protospacers as pTarget that are now flanked by repeat PAMs. (F) In *vivo K*_D_ estimates based on the ratio between Probing/Bound Cascade and the plasmid copy number (Figure S5; Methods). (G) pTarget establishment for *Δcas3* (blue), WT (high induction; green), an empty high copy plasmid (pControl; pink), and low or high copy plasmids carrying CRISPR arrays (pCRISPR2_LC/pCRISPR2; grey/purple). Each dot represents an independent biological replicate. (H) pTarget establishment plotted for different copy numbers of Cascade. Same as Figure 1E but with addition of pCRISPR2. The Cascade copy number of the pCRISPR2 strain was estimated from the relative abundance of the Cascade (probing) fraction in the WT (high induction; Figure 2C) and pCRISPR2 (Figure S4) strain. Each dot represents an independent biological replicate. Error estimation in (A-F) is based on bootstrapping (± standard deviation). See also Figure S3, S4 and S5.

Compared to a non-targeted control plasmid (Figure S3), the introduction of pTarget in cells decreased the fraction of free Cascade complexes (from 60 ± 4 to 29 ± 3%), and gave rise to a 34 ± 2% immobile, target-bound Cascade fraction 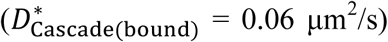 (Figure 5A). As expected, addition of pTarget increased the persistence of sustained binding events, indicating specific DNA target binding (Figure 5C). The combined information of plasmid copy number and the fraction bound Cascade enabled us to determine a cellular *K*_D_ value for the affinity of Cascade for targets of ∼180 nM (Figure 5F; Methods), indicating that the affinity *in vivo* is around 10 times lower than what has been observed *in vitro* (Hayes et al., 2016).

We hypothesized that transcription of DNA along target sites would be one of the main factors influencing Cascade target DNA binding. To investigate the effects of transcription by host RNA polymerase (RNAP), we introduced a (lac) promoter in front of the pTarget sequence. To our surprise, we observed that the affinity of Cascade for target sites that undergo transcription (∼100 nM) was higher than for non-transcribed target sites (∼180 nM). In addition, we observed an increased fraction of free Cas8e subunits (from 37 ± 2% to 54 ± 2%) in the strain containing transcribed pTarget (Figure 5B). Collectively, these findings suggest that transcription of a target DNA sequence somehow facilitates target search and increases the affinity of a target. In addition, it appears that collisions of RNAP with target-bound Cascade result in changes in the Cascade assembly, likely by dissociation of the Cas8e subunit from the complex upon collision with RNA polymerase, which potentially dissociates Cascade from the target.

The relatively dynamic association of Cas8e within the Cascade complex has been observed previously *in vitro* (Jore et al., 2011) and was more recently also observed upon binding to the CRISPR array (Jung et al., 2017). We hypothesized that this dynamic behavior might be a functional characteristic and will also occur upon encountering CRISPR arrays inside the cell. To test this hypothesis, we made a variant of pTarget where all 18 interference PAMs were replaced by the trinucleotide sequence matching the repeats of the CRISPR array (pCRISPR1). Cascade did not show any interaction with the non-transcribed pCRISPR1 plasmid (Figure 5D). However, when we added a promoter sequence in front of the pCRISPR1 array of targets, we observed moderately enhanced levels of free Cas8e (from 40 ± 1 to 56 ± 1%) (Figure 5E), reminiscent of Cas8e expulsion from the complex upon collision with RNA polymerase, or from targets with repeat like PAMs (Jung et al., 2017). Effectively this shows that transcribed CRISPR arrays may function as target decoys in the cell and can therefore potentially influence the levels of functional Cascade complexes in the cell.

To test whether CRISPR array really form decoys in the cell and could impact interference levels, we constructed a compatible high copy number plasmid pCRISPR2 containing a normal CRISPR array (Figure S5). While the introduction of pCRISPR2 into cells containing pTarget only led to a small decrease in the number of Cascade complexes (15% less) (Figure S3), the CRISPR interference levels were reduced by as much as 50% (Figure 5G). This effect was not observed with low copy variant of pCRISPR2 (pCRISPR2_LC) or with a high copy plasmid lacking CRISPR arrays (pControl), indicating that this effect comes from the presence of a large number of CRISPR arrays in the cell (Figure 5G). We further found that the observed impact of CRISPR arrays on Cascade copy number and interference level fits well with our previously predicted relation between Cascade copy numbers and probability of successful MGE establishment (Figure 5H). It furthermore demonstrates how relatively small changes in Cascade copy numbers (15%) can have a big impact on CRISPR interference levels (50%). Taken together, our data indicate that Cascade target search and binding is strongly influenced by the action of RNA polymerase and that CRISPR arrays form target decoys in the cell, which can affect CRISPR interference levels.

## Discussion

How crRNA-effector complexes can achieve timely detection of incoming mobile genetic elements in the crowded environment of the cell is an intriguing aspect of CRISPR biology that remains poorly understood. We provide first insights into the fundamental kinetics of the surveillance behavior of type I crRNA-effector complexes in their native cellular environment. We determined how many copies of Cascade are required to establish effective immunity and uncovered how Cascade complexes navigate the crowded bacterial cell packed with DNA. Our results indicate that Cascade does not restrict its search space to parts of the cell, for example the nucleoid-free periphery, but instead is occupied scanning the entire host nucleoid for a match. To cover this vast sequence space sufficiently fast, the Cascade complex interrogates DNA sequences by using a combination of PAM-dependent and PAM-independent interactions which on average last only 30 ms. This probing interaction is much faster than previously reported interaction times determined of type I Cascade complexes by *in vitro* methods, which range between 0.1 and 10 s (Brown et al., 2018; Redding et al., 2015; Xue et al., 2017). The ability to rapidly probe DNA sequences for potential matches with the crRNA, and to move from one place in the nucleoid to the next, may explain how a relatively low number of Cascade complexes in *E. coli* may still confer CRISPR immunity. Interestingly, the average probing time of 30 ms for Cascade matches values found for *Streptococcus pyogenes* dCas9 in *E. coli* (Jones et al., 2017; Martens et al., 2018), suggesting that DNA probing interactions of crRNA-effector complexes from both Class I and II systems may have evolved independently to take place at this time scale and may.

Given the hundreds of thousands of PAMs in the host DNA, this interaction time would lead to a search time in the order of hours. This value matches our independently calculated estimate of 1.5 hours for a single Cascade to find a single DNA target in the cell, which is four times faster than dCas9 search time estimates of 6 hours (Jones et al., 2017). However, our data also indicates that Cascade not only probes PAMs, the complex also spends a considerable amount of time engaged in PAM-independent DNA interactions. These might be constituted by direct crRNA – DNA interactions (Blosser et al., 2015; Xue et al., 2016), or electrostatic interactions of Cascade with the DNA (Van Erp et al., 2015; Hochstrasser et al., 2014). This suggests an even larger DNA sequence space needs to be covered, creating the need for even more efficient and functionally flexible surveillance solutions. This more flexible probing behavior would be required to recognize targets with mutations in the PAM or protospacer in order to trigger a CRISPR memory update pathway called priming (Datsenko et al., 2012; Jackson et al., 2017), which appears to be unique for type I CRISPR-Cas systems.

One possibility to reconcile Cascade DNA probing characteristics to the overall search time could be that Cascade undergoes facilitated 1D DNA sliding, where Cascade probes multiple sites per DNA binding event. We have shown that Cascade spends 50% of its search time on DNA, and the other 50% diffusing to a new site in the cytoplasm. This value may seem low compared to other DNA interacting proteins such as transcription factor LacI, which is DNA bound for 90% of the time (Elf et al., 2007). However, 50% has been theoretically derived as the optimum for a target search process involving one dimensional DNA sliding and 3D translocation/hopping (Slutsky and Mirny, 2004). Indeed, recently it has been shown *in vitro* that Cascade and Cas9 can slide along the DNA in search of targets (Brown et al., 2018; Globyte et al., 2018). If this also occurs *in vivo*, this would be a striking example of a DNA binding protein having an optimized time division between DNA-bound and freely mobile states to survey the DNA content of the cell.

The relatively high abundance (50%) of freely diffusing Cascade complexes may have benefits as well, as this will lead to more Cascade complexes in the periphery of the cell outside of the nucleoid. By surveying these peripheral regions more frequently, Cascade may be able to detect incoming bacteriophage or plasmid DNA more rapidly when these genetic elements enter the cell.

Besides the chromosomal host DNA, other cellular constituents also affect target DNA binding properties. We found a much higher K_D_ value *in vivo* (180 nM) than was reported earlier using *in vitro* methods (20 nM) (Hayes et al., 2016). The discrepancy in binding affinity between *in vivo* and *in vitro* measurements may be caused by an increase in target search time (i.e. a lower on-rate) or an increase in target dissociation rate (i.e. a higher off-rate) *in vivo*. In any scenario, this discrepancy highlights the strong role of the crowded cellular environment on target binding.

Counterintuitively, we have found that Cascade binds transcribed target sites with higher affinity (100 nM) than non-transcribed target sites (180 nM). Previous studies have shown that negative-supercoiling is required for Cascade binding (Westra et al., 2012), and that increased negative super-coiling accelerates the rate of R-loop formation (Szczelkun et al., 2014). As transcribed regions cause more negative supercoiled regions in the DNA (Ma and Wang, 2016), this could explain the increase in the affinity for transcriptionally active sites. Rates of spacer acquisition were also found to be higher for transcriptionally active regions (Staals et al., 2016), so together these effects may influence the abundance and effectivity of spacers in nature.

Next to the positive effect of transcription on target search, we have also found that collisions between RNAP and target-bound Cascade lead to Cascade disassembly, where the Cas8e subunit is expelled from the Cascade core. Furthermore, also CRISPR arrays themselves can trigger Cascade disassembly, indicating they form target decoys in the cell. When present at high copy number, CRISPR arrays can even impact CRISPR interference levels (Fig. 5G). The loose association of Cas8e with the core Cascade complex as observed *in vitro* (Jore et al., 2011), might serve a biological role in cells to recycle Cascade from off-targets including the CRISPR array, and may prevent Cas3 recruitment and subsequent self-targeting (Xiao et al., 2018).

By measuring cellular copy numbers, and accurately measuring CRISPR interference levels, we could uncover an exponential relationship between the number of Cascade complexes in the cell and CRISPR interference. This relationship describes that every 20 Cascade complexes loaded with one crRNA can provide 50% more protection from an invading DNA element (i.e. 20 copies provide 50%, 40 copies 75% protection). Therefore at constant Cas protein production and degradation levels, the effective concentrations of Cascade complexes loaded with one type of crRNA will become diluted when CRISPR arrays become longer. The size of the CRISPR array is therefore a tradeoff between the higher protection levels of a few spacers, and lower protection levels of many spacers. With our findings we can test optimality of this tradeoff under different conditions and help explain the observed sizes of CRISPR arrays found in nature (Martynov et al., 2017).

The initial entry is the most vulnerable time for the invader, but invading MGEs have the possibility to outrun CRISPR-Cas immunity by replicating faster than being found. In the native cellular environment, we have found that scanning of host DNA, binding to CRISPR arrays and encountering transcribing RNA polymerases can prevent Cascade from finding the target before the critical time (*t*_c_) is reached and the invader is permanently established (Figure 6). We therefore hypothesize the presence of a kinetic arms race, in which invaders have evolved to replicate increasingly fast upon cell entry, while CRISPR-systems have evolved to increase the rate at which they are able to find the target. A recent study has indeed shown that the replication rate of foreign elements affects CRISPR interference levels (Høyland-Kroghsbo et al., 2018). Many bacteriophages use a two-stage injection (Chen et al., 2018; Davison, 2015), which may have evolved to limit the amount of time their DNA is exposed to intracellular defense mechanisms, while already allowing the production of proteins to replicate phage DNA, control host takeover, or to inhibit host defense (e.g. anti-CRISPR proteins) (De Smet et al., 2017). It has been previously shown that the host can counter this strategy by selectively targeting early injected DNA regions, maximizing the time available to look for targets (Modell et al., 2017).

**Figure 6:**
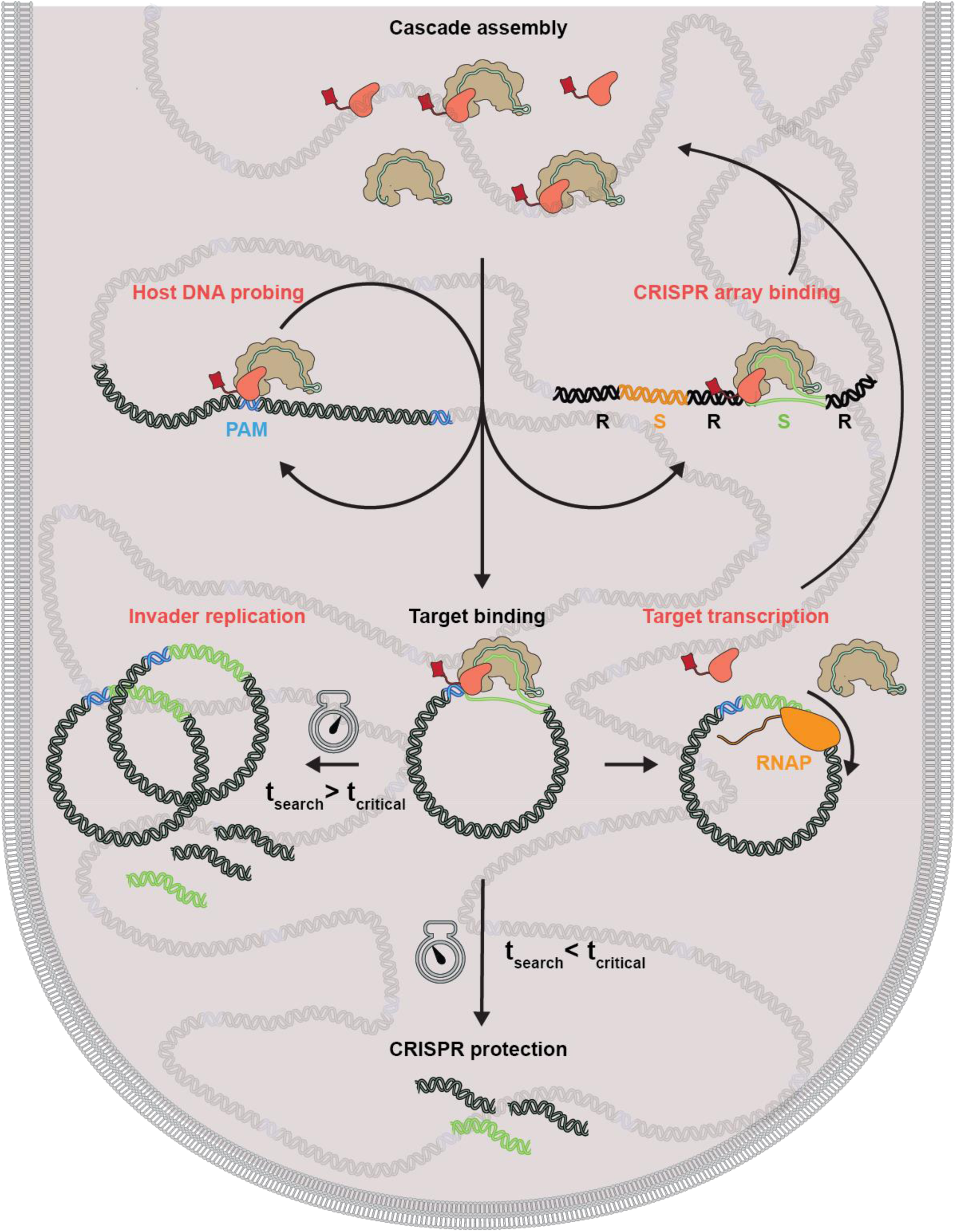
Model of how Cascade protects the cell. Successful protection against an invader requires Cascade target search to circumvent several potential diversions (red). After Cascade is assembled, the complex probes the host DNA by rapidly binding and dissociating. It uses PAM-dependent and PAM-independent DNA interactions and scans the entire nucleoid region. If it binds to a CRISPR array (S: spacer; R: Repeat), the complex disintegrates. When it has found its target, it depends on the search time (*t*_*search*_) and the critical time (*t*_*critical*_) whether the invader is cleared and the cell protected, or the invader can replicate and establish itself in the cell. Moreover, transcription by RNA polymerase (RNAP) can still remove bound complexes, compromising CRISPR protection.

Our mathematical description of CRISPR interference can be adapted to natural environments in which the growth rates are orders of magnitude slower than under laboratory conditions. Furthermore, the target search equations established here could be expanded to the population level, allowing to model how individual variability in Cascade expression levels and growth rates can impact the survival of entire populations. Therefore, our data provides an important framework for further quantitative cellular studies that will address how CRISPR systems optimally deal with the challenges of cost-effective and rapid target search.

## Methods

### Cloning

The inserts to create pTarget and pCRISPR1 plasmids were purchased as synthetic constructs from Gen9 (pTarget insert and pCRISPR1 insert; Table S3). To increase the copy number of targets in the cell, the constructs were cloned into a pUC19 backbone with XbaI and KpnI restriction sites, yielding pTarget-RNAP and pCRISPR1-RNAP. The lac promoter was removed for both plasmids by digestion with SalI and PciI, creating blunt ends with Klenow Fragment and subsequently religated to yield pTarget and pCRISPR1. CRISPR arrays were amplified from the K12 BW25113 strain (primers BN383 and BN384; BN370 and BN385 for CRISPR array 2.1 and 2.3 respectively) and cloned into pJPC-12 plasmid containing the pSC101 ori with KpnI and SalI sites (for CRISPR array 2.1) and SalI and EcoRV sites (for CRISPR array 2.3). The copy number of the plasmid could be varied by introducing mutations in the *repA* gene with site-directed mutagenesis PCR (BN373-375). The E96R mutation of RepA yields a reported copy number of ∼240/cell (pCRISPR2) compared to the WT RepA (pCRISPR2_LC) copy numbers of ∼7/cell (Peterson and Phillips, 2008). A plasmid was made from the high copy-variant that did not contain any CRISPR arrays (pControl). All constructs were verified by sequencing.

### Recombination

The strains used in this study were created by using Lambda red recombineering (Datsenko and Wanner, 2000). Strains harboring the pSC020 plasmid that contains both the Lambda red recombinase and Cre-recombinase were grown at 30 °C. Before transformation of an insert containing an antibiotic resistance marker, the expression of Red recombinase was induced with 0.2% L-Arabinose. Colonies on the specific antibiotic plate were verified with PCR and sequencing and subsequently Cre recombinase expression was induced with 1 mM IPTG at 37 °C to promote plasmid and antibiotic resistance gene loss. The strain was subsequently patch plated to screen for resistance sensitivity due to plasmid loss.

If the scar that is left after lox-site recombination is directly upstream or downstream of a gene it might influence gene transcription/termination. In the design of constructs for *pamcherry2* (Subach et al., 2009) the lox*-cat-*lox sequence was placed upstream of the IGR (Intergenic region) that is present between *cas3* and *cas8e*. To allow for correct termination of *cas3,* a part of the IGR was also added at the 5’ end of the antibiotic resistance marker. The 3’ flank of the constructs overlapped with the *cas8e* gene. The 5’ flank of the constructs matched a sequence upstream and downstream of *cas3* (PAmCherry ins; Table S3). Amplification of the constructs with a forward primer matching the downstream region kept *cas3* intact upon insertion (BG7128), whereas a primer matching the upstream region deleted the *cas3* gene allowing measurements in the presence of targets (BG7129). The insert also contained a part of the *cas8e* sequence containing a G160A mutation. This mutation could be introduced into the gene simultaneously with the fluorescent protein, depending on the reverse primer that was used for insert amplification (BG7130 for WT, BG7131 for G160A).

Knockouts of the CRISPR arrays and Cas gene subunits of the K12 strain were made by amplifying a lox-*kan*-lox or lox-*cat*-lox sequence with flanks matching the specific sequences and introducing them into the strain as described above (BG7366+BG7367 for CRISPR array 2.1; BG7368+BG7369 for CRISPR array 2.2+2.3; BG8366+BG8367 for Δ(*cas11-cas6e*)). A full overview of the sequences of these inserts is given in Table S3.

### Growth conditions

To prevent the high-copy target plasmids from influencing the growth rate of the strains and therefore changing the fraction of matured PAmCherry complexes we used a rich defined medium with minimal autofluorescence. Strains were grown in M9 minimal medium containing the following supplements: 0.4% glucose, 1x EZ amino acids supplements (M2104 Teknova), 20 μg/ml uracil (Sigma-Aldrich), 1mM MgSO_4_ (Sigma-Aldrich) and 0.1 mM CaCl_2_ (Sigma-Aldrich) (further referred to as M9 medium). Strains were inoculated o/n from glycerol stocks and 200x diluted in fresh medium the next day. Cells were always grown with the required antibiotics. The expression level of Cascade for strains carrying the pKEDR13 plasmid could be tuned by different expression levels of LeuO. The expression level referred to in the text as low induction was achieved by leaky expression of LeuO (no addition of IPTG), whereas high induction was achieved by addition of 1 mM IPTG upon dilution of the o/n culture. For all sptPALM measurements the high induction condition was used. The cells were grown for ∼2.5 hours to an OD of 0.1 before use. For enforced elongation of cells, cephalexin (40 μg/ml) was added 0.5 hour after fresh inoculation and grown for two more hours. When required, DAPI for staining of DNA was added right before imaging (0.5 mg/ml).

### Transformation assay

Each culture was grown under conditions described above and 30 ml were used to create competent cells. Cells were washed 3 times in ice-cold 10% glycerol solution and the final culture was reduced to 250 µl. The cells were aliquoted and stored at −80 °C. A mixture of pTarget (10 pg/μl) and pGFPuv (10pg/μl) was transformed into 40 µl of culture. In case of strong interference levels, the ratio was adjusted to a 100:1 (pTarget (100 pg/μl):pGFPuv (1 pg/μl)). The transformability of strains was linear in these concentration regimes, allowing these different relative concentrations to be used.

Electroporated cells were immediately plated in two dilutions on plates containing ampicillin (100 µg/ml) and glucose (0.4%). Glucose was added to prevent premature expression of GFPuv which would cause a decrease in fitness of cells containing this plasmid. The next day, 96 colonies from each replicate were reinoculated in 96-wells plate with LB containing ampicillin (100 µg/ml) and IPTG (1 mM). After overnight incubation, the 96 well colonies were analysed in a plate-reader (Synergy H1, Biotek). pTarget establishment was defined as

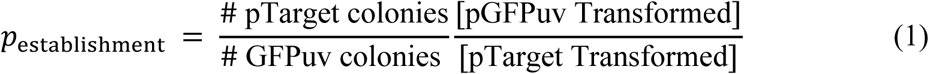

pTarget establishment was further normalized to the interference level of a Δ*cas3* strain.

### qPCR

Each culture grew under conditions described above and 2 ml were used to extract the DNA. DNA was isolated with the Genejet Genomic DNA kit (Thermo Scientific) and concentrations were measured with the Qubit dsDNA HS Assay kit (Thermo Scientific). qPCR was performed with primers that have been used before in plasmid copy determination (BG8677-BG8680) (Reyes-Lamothe et al., 2014). The Ct value of the PCR amplifying the *dxs* gene and the *bla* gene was a measure for the ratio between chromosomal and plasmid DNA. 1 ng of genomic DNA and 0.5 μM of each primer was added to the iTaq^TM^ SYBR Green SYBR Green PCR reaction mixture. A standard curve for the amplification efficiency was made by a dilution series of pMS011, a plasmid containing one copy of the *dxs* and the *bla* gene.

### Slide preparation

In order to work with very clean slides, an extensive cleaning procedure was used (modified from (Chandradoss et al., 2014)). Slides were burned in the oven at 500 °C for two hours, and stored in aluminum foil until the day of usage. Slides were subsequently sonicated in MilliQ, Acetone and KOH, incubated in Piranha Solution (75% H_2_SO_4_, 7.5% H_2_O_2_) and afterwards rinsed with MilliQ. 1% Agarose slabs containing the growth medium were hardened between two cleaned glass slides, spaced slightly apart using parafilm. After hardening, a concentrated culture of cells was added in between the slab and one of the slides. The agarose slab was always prepared within 20 minutes of the measurement to prevent desiccation.

### Microscope set-up

For the acquisition of microscopy data, a home-build TIRF microscope was used, which is described in more detail elsewhere (Martens et al., 2018). Briefly, four lasers with different wavelengths (405, 473, 561 and 642 nm) are situated in a Lighthub laser box (Omicron, Germany), and are transformed in a collimated beam via a reflective collimator and an optical fibre. Stroboscopic illumination was used to allow for 2 ms excitation in the temporal middle of the captured 10 ms long frame (Farooq and Hohlbein, 2015). The excitation laser is focused on the backfocal plane of a 100x oil immersion SR/HP objective (NA = 1.49, Nikon, Japan), and the emission is captured on a Zyla 4.2 plus sCMOS camera (Andor, UK). 2×2 pixel binning was used, resulting in 128×128 nm pixels. Data acquisition was performed using MicroManager (Edelstein et al., 2010). Measurements were performed at room temperature (21 ° C)

### Single-molecule Measurements

The cells were imaged with a brightfield light and 405 and 561 nm lasers. First brightfield images were taken to find contours of the cells. The 405 nm laser was used to stochastically activate PAmCherry and the laser intensity was slowly increased during the measurement up to 10 μW. The laser intensities were measured directly after the reflective collimator. With increasing the laser intensity of the 405 nm laser during the measurements, we aimed at keeping the number of activated molecules relatively constant (∼1-10 per FOV). The 561 nm laser was used to excite the fluorescent protein tags (40 mW pulses with 2ms pulse width, leading to average exposure intensity of 8 mW).

To measure Cascade localization in cephalexin-treated cells that were stained with DAPI, we took an alternative approach. To prevent DAPI fluorescence from influencing the fluorescence measurements of the single molecules, we briefly activated a subset of particles with the 405 nm laser and subsequently tracked Cascade for a couple of frames with 561 nm excitation, repeatedly doing this, until most fluorescent proteins were photobleached.

## Analysis

### Detection, localization and tracking

Analysis was done with home-built software, adapted from (Holden et al., 2010; Uphoff et al., 2013). The sCMOS camera we used has pixel dependent offset, gain and variance, which we took into account to minimize the detection of false positive localisations. We estimated these parameters by measuring 60.000 dark frames and 20.000 homogeneously illuminated frames with increasing levels of intensity (Vliet et al., 1998). To further optimize our detection, we implemented a temporal median filter (time window 400 frames) for background estimation (Hoogendoorn et al., 2015). The background estimate was not directly subtracted from the image, but photon statistics were incorporated in a likelihood-ratio test that calculated the probability of a scenario with and without an emitter for each pixel in every frame. Briefly, a raw image was first converted into photon counts by using the camera offset and gain maps. Subsequently for every pixel the intensity (*I*_tot_) of a potential emitter was estimated by Gaussian-weighted (σ =1 pixel) summation of a 7×7 window to a background subtracted image. Subsequently, potential emitters of more than 50 photons were preselected and were further subjected to a ratio test. The ratio test uses the probability defined for pixel *i* to have a transformed value *v* in the 7×7 region around the preselected pixels as previously described (Huang et al., 2013):

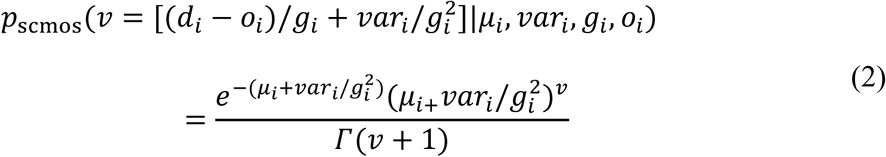

Where *d*_i_ is the raw image value, *g*_i_ is the gain, *var*_i_ the variance and *o*_i_ the offset for pixel *i*. The ratio test calculates the product of the probability of all pixels in the subregion in case of an emitter *μ*_*i*_ = *b*_*i*_ + *I*_*i*_, where *b*_i_ is the estimated background an *I*_i_ is the estimated intensity of the emitter at pixel *i* (which was estimated by a Gaussian from the center of the 7 × 7 subregion with emitter intensity *I*_tot_) divided by the product of the probability of all pixels in the subregion in case of absence of an emitter *μ*_*i*_ = *b*_*i*_.

We set the likelihood to a level that achieved approximately one false positive per frame of 512 × 512 pixels. This method allowed the detection efficiency to be more robust across and between FOVs and independent of manual thresholding for each measurement. Detected particles were subsequently localized with MLE-sCMOS software as previously described (Huang et al., 2013).

The localized particles were subsequently linked. Localizations in subsequent frames that were closer to each other than 6 pixels in length (0.78 μm) were assigned as a track. Particles were allowed to disappear for one frame (due to blinking/moving out of focus), but these steps were not used in the calculation of the apparent diffusion coefficient, *D*\*.

### Determination of diffusion coefficients

Several methods were employed to extract diffusion states and their abundances from the analysed tracks. The distribution of the apparent diffusion coefficients can be fitted to an analytical equation as reported earlier (Stracy et al., 2015; Vrljic et al., 2002). These equations depend on the number of steps that is used to generate the average diffusion coefficients of each particle. We used tracks containing a minimum of four steps and only four steps were used in longer tracks.

For a single diffusion coefficient fitting becomes:

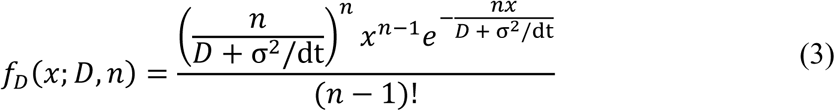

With multiple states this equation becomes:

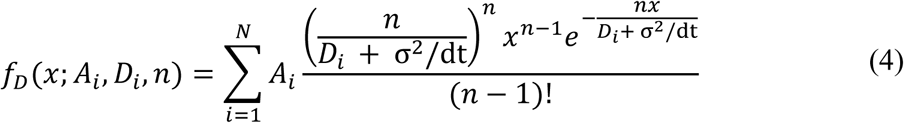

Where *A*_i_ are the fractions (∑ *A*_*i*_ = 1), *D*_i_* are the apparent diffusion coefficients of the different states and *n* are the number of steps. The localization error (σ) was found to be 40 nm, based on the apparent diffusion of the slowest moving fraction in our global data set and similar to other studies using the same fluorescent protein (Stracy et al., 2015; Uphoff et al., 2013) or set-up (Martens et al., 2018). This equation was fitted to our track distributions with a Maximum Likelihood Estimation algorithm. The uncertainty in the fit was estimated with Bootstrap resampling. The list of *D*\* values was resampled 20.000 times with replacement to the size of the original data set. Each resample was then fitted with the same Maximum Likelihood Estimation algorithm.

### Analytical Diffusion Distribution Analysis (DDA)

*D*\* Distributions have been fitted in numerous studies of DNA binding proteins (see above) (Stracy et al., 2015; Vrljic et al., 2002), making use of distributions developed by Qian *et al.* (Qian et al., 1991). The goal is to find the distribution of measured *D*\* values (*x*), for a certain number of underlying states that each have a probability *A*_i_ and a diffusion coefficient *D*_i_. It is derived from repeated convolution of the exponential distribution of displacement, resulting in a gamma function for each state. These distributions assume, however, that there is no transitioning occurring between states.

In order to incorporate dynamics of state transitions into our fitting, we incorporated statistics coming from photon distribution analysis (PDA) that is used for single molecule FRET diffusion coefficient distributions (Antonik et al., 2006; Kalinin et al., 2008; Palo et al., 2006). This method, that we term Diffusion Distribution Analysis (DDA), describes the distribution of time spent in each state given a certain, 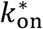, *k*_off_ and the integrated time *t*_int_. Here we discuss the analytical way to find this distribution.

Firstly, the probability distribution function for time can be calculated by three equations corresponding to 0, an odd and an even number of transitions (Palo et al., 2006):

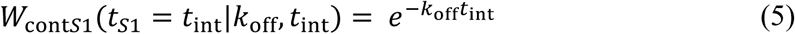

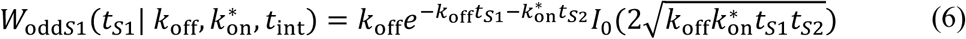

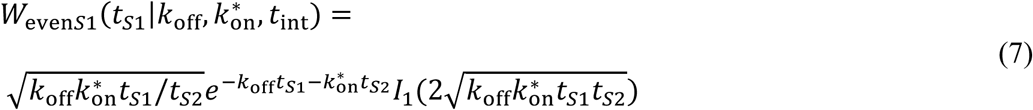

Where *t*_*S*1_ and *t*_*S*2_ are times spent in state *S1* and state *S2* and *I*_0_ and *I*_1_ are Bessel functions of order zero and one respectively. Note that *t*_*S*1_ + *t*_*S*2_ = *t*_int_. Equations for starting in state 2 (W_contS2_, W_oddS2_ and W_evenS2_), can be found by exchanging *k*_off_ for 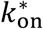 and *t*_S1_ for *t*_S2_ and vice versa in equations 5-7.

We can convert the time spent in the mobile state (*t*_*S*2_) to the diffusion coefficient by the following equation:

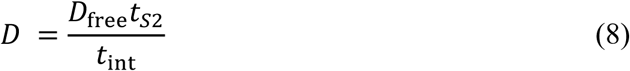

It follows that the probability distribution functions can be converted by:

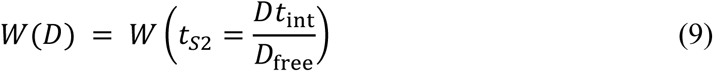

Furthermore, the chance that the particle at the start is in state 1 or state 2 is provided by:

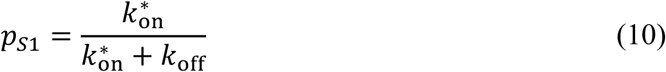

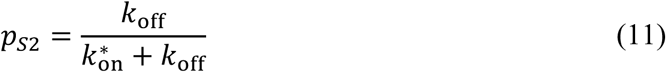

To correctly describe the distribution over a certain number of frames, we first calculated the distribution over a single time frame *t*_*f*_. Within a single frame, a particle started in that state can either end in the same state or in a different state. Therefore, in a two-state system the probability function for four scenarios have to be calculated:

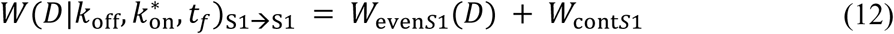

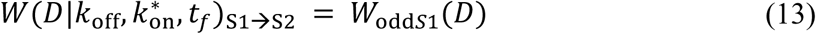

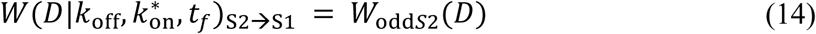

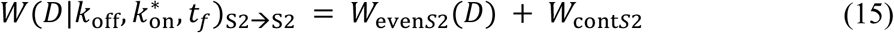

Subsequently the probability to find a certain diffusion coefficient (*x*) for a single time step given the underlying average diffusion coefficient (*D*) is given by *f*_*D*_(*x*|*D*, 1) (Eq. 3). Then we find the distribution of measured diffusion coefficients for a single frame by:

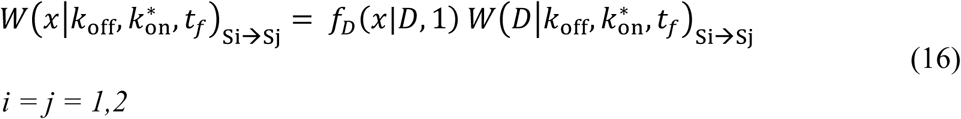

Now that we have the distribution for a single time step, we need to find the distribution for the average of multiple frames. For this we use the same method as Qian *et al.* (Qian et al., 1991), namely repeated convolution of the distribution for a single frame, while keeping track of the start and end state. The probability distributions are therefore:

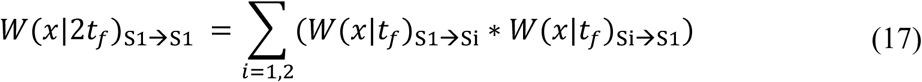

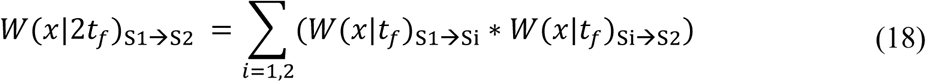

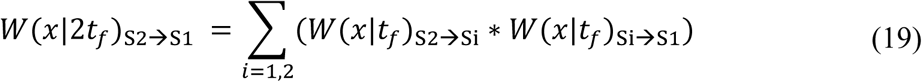

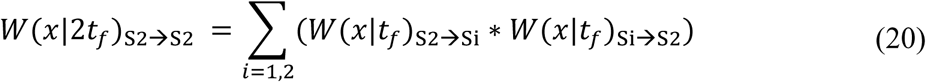

For 4 frames, the distributions found for 2 frames can be convoluted again. The full distribution is then found by summing up each of the partial distributions multiplied by the chance they start in *S1* or *S2*:

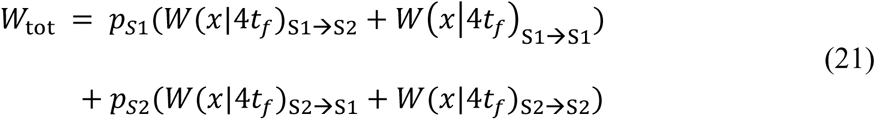

We then have to further correct for the broadening of the distribution of immobile particles where the apparent step size comes from localization error (Figure S3). As localization error, in contrast to diffusion, is correlated (Michalet, 2010), the distribution is not described by a gamma distribution, or any other known exact solution. We find very close agreement with simulations when we subtract the fraction of immobile particles after four time steps (*W*_*contS*1_(*t*_*S*1_ = 4*t*_*f*_), Eq.5) multiplied with the distribution of expected *D*\* for four time steps *f*_*D*_(*x*|0,4) (Eq. 3) and replace it with the same fraction of immobilized particles multiplied with the distribution of expected *D*\* for 2.9 time steps *f*_*D*_(*x*|0,2.9). This value stems from the variance found for correlated MSD values due to localization error (Michalet, 2010).

### Copy number determination

The copy number of the Cascade complex was determined by generating cell outlines from brightfield images (only well separated cells were chosen). The cell outlines were made with the Oufti software (Paintdakhi et al., 2016). The total number of tracks that were found in the outlined cells generated a copy number (Figure 1D). Because single localization events can partly stem from false positives, the total amount of tracks was estimated based on the distribution of tracks longer than 1 step and subsequently this distribution was fitted with an exponential to calculate the amount of particles that only had a single localization before bleaching. Similarly, as we know the false positive rate was approximately one per frame, we could also subtract the number of frames from the single step tracks and in this way estimate the total number of tracks. This approach yielded comparable results.

The copy number of proteins in cells are hard to quantify (Lee et al., 2012). Currently, protein copy numbers can be estimated either by western blot or by single-molecule fluorescence based methods both of which have specific drawbacks. Although single molecule studies are regarded as the most accurate method, especially at low copy numbers (Huang et al., 2007), there are a lot of variables that can lead to over- or underestimation. Underestimation can originate from maturation time of the protein, misfolded/inactivated protein, false negative detections, overlap of PSFs and linking of two separate molecules in a single track. Overestimation can come from failed linking of tracks, false positive detections and blinking fluorescent proteins.

As has been done in previous studies, we take the underestimations stemming from maturation time (23 min for PAmCherry (Subach et al., 2009)), close to growth rate of 31 min) and estimated *in vivo* folding efficiency (50% (Durisic et al., 2014)) into account (Uphoff et al., 2013). We also consider that an estimated 40% of the particles we observed come from Cas8e subunits not active complexes. Taken together, the number of particles we observe are subtracted by the amount of estimated autofluorescent particles and subsequently multiplied by a compensation factor of two to reach our estimated copy number values.

We believe that the assumptions made in this study could maximally lead to over- or underestimating our estimated copy numbers by two to three-fold. We note that the relative amounts we observed between the different expression levels will be independent of these assumptions.

### Cascade in DNA-containing/DNA-free regions

To get an independent measure of the total time fraction spent probing DNA, Cascade was visualized in cells that were elongated by addition of cephalexin. The drug cephalexin disabled the ability of the cells to divide, creating elongated cells where nucleoids were separated by DNA-free spaces (Reyes-Lamothe et al., 2014). Subregions of cell outlines were manually selected and further refined with the Oufti software (Paintdakhi et al., 2016). The relative amount of localizations of DNA-free and DNA-containing regions was not calculated for entire cells, as differences in illumination intensity between parts of the FOV could also change the amount of localizations detected for different parts of the cell. Each subregion contained one nucleoid free region, flanked by two nucleoid containing regions with a total length of around 4 μm. Segments of 0.1 μm divided along the long axis of the cell are separated into nucleoid or DNA-free segments based on the sum of the DAPI fluorescence within each segment. The average number of localizations of Cascade molecules in nucleoid segments divided by the average number of localizations Cascade molecules in DNA-free segments could be used to infer the DNA bound time fraction (see below, f_onDNA_ from nucleoid enrichment).

### Persistence sustained binding events for different integrated times

To estimate how long binding events last, one could plot the number of particles remaining within a certain radius from the first frame position for different number of steps. However, particles can diffuse away when they are released from DNA or be lost due to photobleaching. To account for bleaching rates, previous studies increased dark time between exposures, while keeping exposure times the same (Ho et al., 2018; Knight et al., 2015). This approach uses the data of all time steps, including only single time steps.

As we are investigating lifetime of binding events on a subsecond timescale this approach fails, as single steps of slow-moving particles, which can be clearly separated from bound particles on larger timescales (*t*_int_ > 1 s), will be counted as bound particles leading to overestimated off-rates. At these timescales, it is more reliable to use tracks of at least 5 steps to distinguish bound from moving particles. As we are interested in how many of these events we observe, depending on the framerate, normalization is required.

For this we cannot use the sum of all tracks observed at each frametime, as a larger amount of fast moving molecules diffuse further than the maximum tracking distance of 0.78 μm between two exposures, and are also more affected by confinement with increasing integrated time. Therefore, the number of moving particles of certain track length is not an accurate normalization when comparing different frame times. However, as we used similar exposure for all frame times, the number of detected localizations per protein is unaffected. Furthermore, bound molecules are not affected by confinement or linking errors with increasing frame rates.

The most robust normalization procedure was therefore to normalize the number of localizations within sustained bound tracks (all localizations within 1 pixel of the mean location of the track) to the total number of localizations, as those do not depend on the length of introduced dark time between exposures. A further increase of the dark time was not possible as on longer time scales the movement of the plasmid 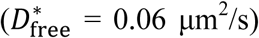 made plasmid bound particles diffuse further than 1 pixel.

### Confinement and localization error simulation

To verify whether our new transitional *D*\* analysis yielded accurate parameter predictions and investigate the influence of localization error and confinement on the parameters of the fit, we simulated particles moving and transitioning between bound and free moving states within the dimensions of an *E. coli* cell, adapted from methodology used in (34). At every time step particles were simulated to be either in a bound state *S1* (*D* = 0 μm^2^/s), or a mobile state *S2* (*D* = *D*_free_). At the starting time point, states were assigned to each particle according to the equilibrium probability *p*_*S*1_ and *p*_*S*2_ (Eq. 10 + 11). Subsequently, at following time steps of 0.1 ms, particles in state *S1* were assigned to *S2* with a probability of *p*_*S*1→*S*2_ = *k*_off_*t*_step_ (where *t*_step_ = 0.0001 *s*) and particles in state *S2* were assigned to *S1* with a probability of 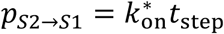. Displacements in three dimensions at each time step were taken from a standard normal distribution multiplied with 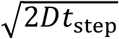 (where *D* is either 0 for particles in state *S1* or *D*_free_ for particles in state *S2*). Steps beyond the boundaries of a cell were rejected and new displacements were randomly drawn.

The 2D projection of five localizations at 10 ms time intervals for each molecule was generated as output and was analysed in our tracking software. Localization error was included in the simulation by addition of a random displacement for each position taken from a Gaussian distribution (*σ* = 40 nm). It was found that changes in outcome of the simulation were not sensitive to cell length in the range of our bacteria (3-6 μm), decreasing less than 5% for the smallest size. Most of the confinement effect is caused by the cell width, which was relatively constant between all the cells measured.

### Cascade nucleoid enrichment simulation

The simulation above was adapted to simulate the movement in DNA-free and DNA-containing regions. Particles were simulated to move inside of a cell of 10 μm in length and 1 μm in width consisting of 100 segments without endcaps (0.1 μm per segment). Five segments were modelled as DNA-free segments and the rest of the segments as DNA-containing segments.

Cascade molecules were randomly placed throughout the cell and subsequently were simulating with similar time steps as described above, except that moving particles were only allowed to transition to *S*1 (bound state) inside of the nucleoid containing regions. Before recording the position of the simulated particles, the simulation ran for 100.000 time steps (10 s) so that equilibrium was reached. Localization error was added in the same way as described above.

### Expected free diffusion coefficients

The diffusion coefficient of molecules in classic (Newtonian) fluids can generally be estimated by the Stokes-Einstein equation. A study measuring the diffusion of GFP multimers inside the *E. coli* cytoplasm has shown good agreement with the predictions of this equation (Nenninger et al., 2010), whereas a second study found a different relation attributed to the complex nature of the cytoplasmic fluid (Mika and Poolman, 2011). To compare our findings of the apparent free diffusion coefficient of Cas8e (∼3.5 μm^2^/s) and Cascade (∼ 1.0 μm^2^/s), we therefore looked for reported free cytoplasmic diffusion coefficient values of proteins of similar size inside *E. coli* cells. For Cas8e, two proteins have been studied with a similar size to PamCherry-Cas8e (82 kDa), namely CFP-CheR-YFP (86 kDa) (Kumar et al., 2010) and TorA-GFP3 (84 kDa) (Nenninger et al., 2010), which have reported values of 1.7 μm^2^/s and 6 μm^2^/s. Our estimate for Cas8e lies within the range of these values. For Cascade (430 kDa), the closest reported protein in size is RNA polymerase, for which the *D*^*\**^_free_ was found to be 1.1 μm^2^/s (400 kDa core enzyme, 470 kDa holoenzyme) (Stracy et al., 2015). Furthermore larger proteins such β-Gal-GFP_4_ (582 kDa; 0.6 μm^2^/s) (Mika et al., 2010), and 30S ribosome subunits (900 kDa 0.4 μm^2^/s) (Sanamrad et al., 2014) were reported with lower diffusion coefficients as expected. These findings support the free apparent diffusion value we found for Cascade (∼ 1.0 μm^2^/s).

### *f*_onDNA_ from nucleoid enrichment

The distribution of Cascade in nucleoid-free and nucleoid containing regions depends on the time Cascade spends on DNA. We divided the cell up along the long axis into segments of 100 nm wide. During the time Cascade is bound to DNA it can only be inside of the nucleoid regions whereas, when it is not bound to DNA Cascade can be anywhere within the cell. Therefore, the average number of particles in a DNA-containing segment is given by:

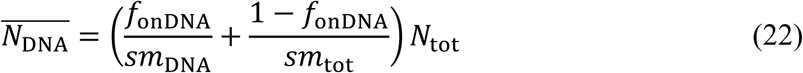

and the average number of particles in a DNA-free segment is given by

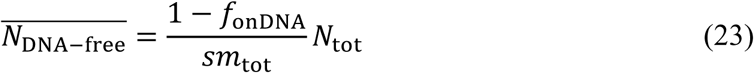

Where *f*_onDNA_ is the fraction of time bound to DNA, *sm*_DNA_ and *sm*_tot_ are the number of DNA segments and the total number of segments respectively and *N*_tot_ is the total number of particles in a cell. The ratio, which is equal to the enrichment factor *EF*, can then be expressed as:

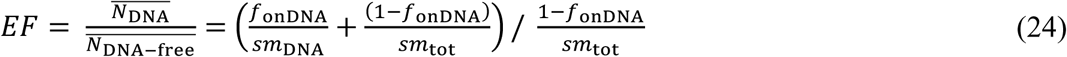

If the number of DNA-free segments is much less than the number of DNA segments *sm*_DNA_ ≈ *sm*_tot_ the expression above can be simplified to:

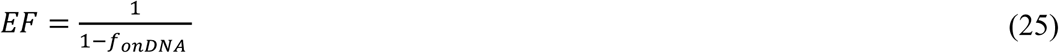

This equation allows extraction of *f*_onDNA_ from EF directly and implies that this value does not depend on the diffusion coefficients of the mobile population.

### *In vivo K*_D_ values

The *K*_D_ value is a commonly calculated affinity constant used for binding kinetics of proteins and assembly of multicomponent systems (McGuigan et al., 2006), but the *K*_D_ has also been used as an estimate for in vivo binding affinity (Zawadzki et al., 2015). In the reaction scheme

A + B ⇄ AB, the *K*_D_ is calculated as

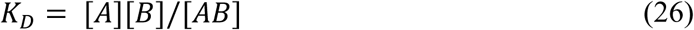

For Cascade the reaction scheme is as follows: [Cascade (probing)] + [free target sites] ⇄ [Cascade (bound)]. The concentration of a single entity inside of a cell of length 4 μm and width 1 μm with hemispherical endcaps is approximately 0.5 nM. The copy number for pTarget was estimated by qPCR to be approximately 100 plasmids per chromosome. As the number of chromosomes in actively dividing cells is generally higher than one, we used literature values for the number of chromosomes/cell found in (Wallden et al., 2016), providing 4/cell which also used a glucose and amino acid enriched M9 medium as growth medium. This brings the copy number of pTarget to 400/cell, which is equal to 200 nM. For a Cascade complex carrying one of several crRNAs in the cell, the amount of free target sites is equal to the copy number of the plasmid pTarget minus the amount of already occupied target sites of that crRNA, but as the copy number of each target (400) is much higher than the number of Cascade complexes potentially carrying that crRNA (on average 130/18 ≈ 7), [free targets] ≈ [pTarget].The *K*_D_ value was then calculated as:

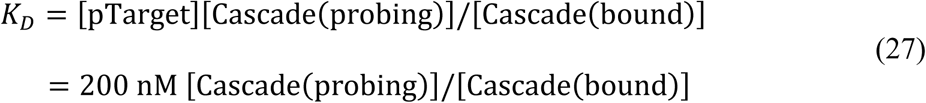

### Theoretical model interference level vs copy number

In the case where the interference level is limited by the target search of the proteins, we can model the relation based on the distribution of search times of single proteins. The search time for each Cascade protein individually is exponentially distributed:

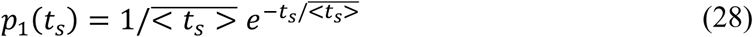

The chance that one of *n* proteins finds the target at search time *t*_s_ while the other proteins have not yet found the target is:

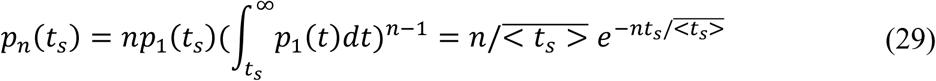

The establishment probability of the plasmid is equal to the likelihood for all search times larger than *t*_critical_ (*t*_c_), the time point at which the cell can no longer clear the invader. Therefore:

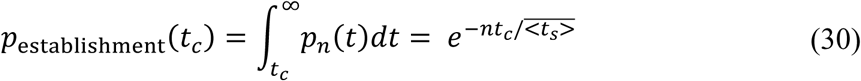

As the chance of targeting after replication is low, we assume in our model that Cascade is only able to clear the foreign DNA before replication. Therefore *t*_*c*_ is equal to the replication time of the plasmid *t*_*R*_.

As we found that 20 copies of Cascade reduce interference level by half, this leads to

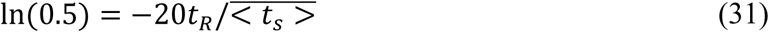

Or

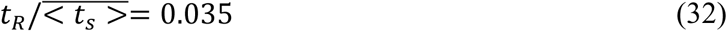

Right after transformation, the negative regulators of copy numbers are absent, so replication in that instant is faster than the growth rate of the cell. Replication time of pTarget has not been measured so far, but by using a temperature-dependent ori, Olsson *et al.* measured a replication time of 3 min for a slightly larger plasmid in the absence of copy number control (Olsson et al., 2003b). If we assume pTarget replication occurs on a similar time scale, we get an estimated search time for one Cascade to find a single target of ∼90 minutes.

## Data availability

The data that support the findings of this study are available within the paper.

## Supplementary information

Supplementary information is available

## Acknowledgements

The authors thank Dr. A. Košmrlj (Princeton University) for deriving equation 29 presented in Methods. We acknowledge S. Creutzberg for supplying plasmid pSC020 and M. Siliakus for supplying plasmid pMS011 and all members of the Hohlbein and the Brouns groups for input during group discussions. We thank Jaap Keijsers, Fiona Murphy and Stan van de Wall for providing preliminary measurements and scripts for data analysis. S.B. is supported by the European Research Council (ERC) Stg grant 639707, and by a Vici grant of the Netherlands Organisation for Scientific Research (NWO). R.M is supported by the Frontiers of Nanoscience (NanoFront) program from NWO/OCW.

## Author contributions

S.B. and J.H. conceived and supervised the project; J.V., M.V. R.M., C.A., D.B., B.B. did the experimental work; J.V. and J.H. derived the theory; J.V. and K.M. wrote analysis scripts; J.V., K.M. and J.H. established microscopy workflow, J.V., J.H. and S.B. wrote the manuscript with input from all authors.

## Author information

The authors declare no competing financial interests. Correspondence and requests for materials should be addressed to S.B. or J.H.

## Supplementary figures

**Figure S1.**
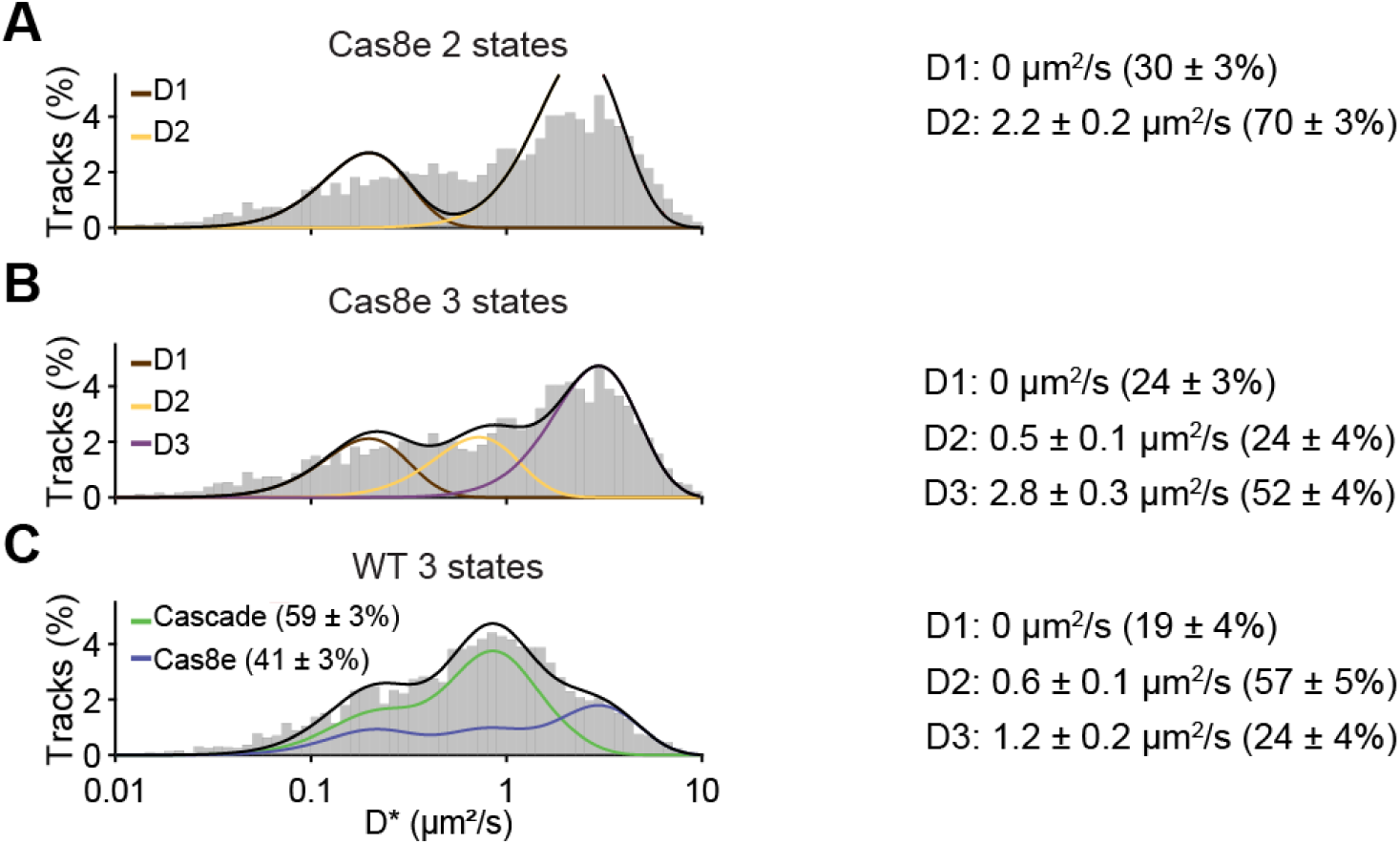
Static *D*\* fitting. **(A)** *D*\* distribution (left) of the Cas8e strain (Figure 2B) fitted with two static states with extracted *D*\* value of each fraction on the right (relative abundance). The slowest state (D1; brown) was fixed to 0 μm^2^/s. **(B)** Same as (A) but then for three static states. **(C)** *D*\* distribution (left) of the WT strain (Figure 2C). Cas8e distribution from Figure S1B was taken and used to fit the distribution with additional three states for Cascade diffusion. The relative abundance of Cas8e and Cascade estimated from static *D*\* fitting is similar to that found for dynamic fitting (60 and 40%), even though the distributions of Cascade and Cas8e are different. Error estimation in (A-C) is based on bootstrapping (± standard deviation).

**Figure S2:**
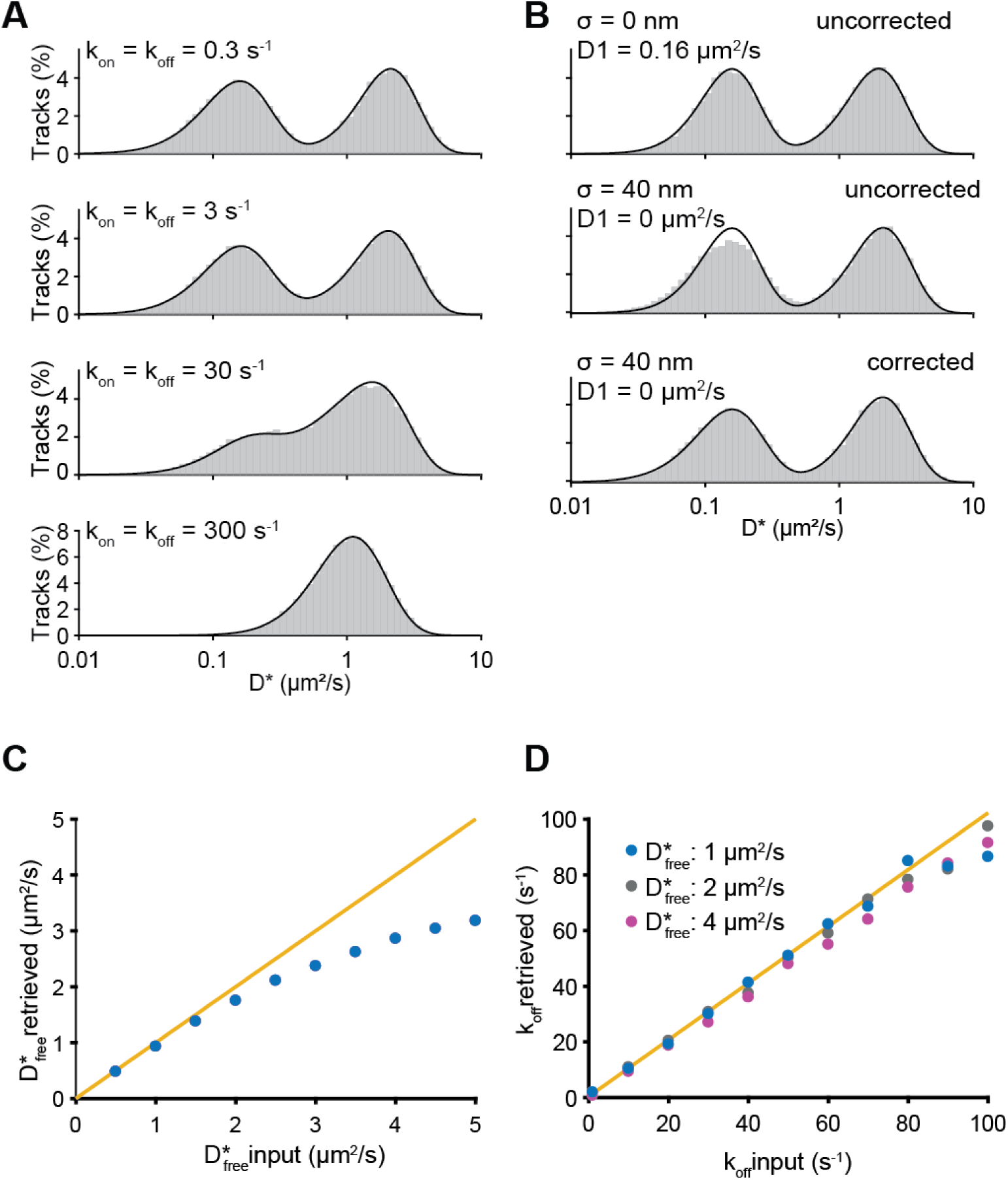
Performance of analytical DDA. (**A**) Comparison of simulation to the theoretical distribution (black line) found with the newly developed analysis method. 50.000 particles were simulated to move without boundaries and position was recorded for 4 consecutive steps. Particles were simulated with 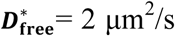 and increasing on- and off-rates (from 0.3 to 300 s^-1^). The theoretical model (black line) is directly plotted on top of the histogram of simulated *D*\* values. A localization error drawn from a Gaussian distribution with σ = 40 nm was added to both the model and the simulation. (**B**) Influence of localization error. Distribution of an average of consecutive displacements that are offset by a localization error are correlated, which is why in the absence of localization error in the simulation (top) there is no requirement for correction. However immobile particles offset by localization error with the same mean apparent diffusion coefficient are slightly differently distributed (middle). Correction (described in Me) for the immobile particles is sufficient to restore the fit (bottom). (**C**) Influence of confinement. Particles were simulated inside of a cell 4 μm long and of 1 μm diameter. Simulations were run through analysis software to retrieve parameters. ***D*^*\**^_free_** estimates are influenced by confinement where fast moving particles appear to be slower. (**D**) The off-rate is not as influenced by effects of confinement and stays the same even for the fastest moving particles (purple). Estimates become more unreliable for much faster or slower transitions than are measured in the integrated time of typical tracks.

**Figure S3.**
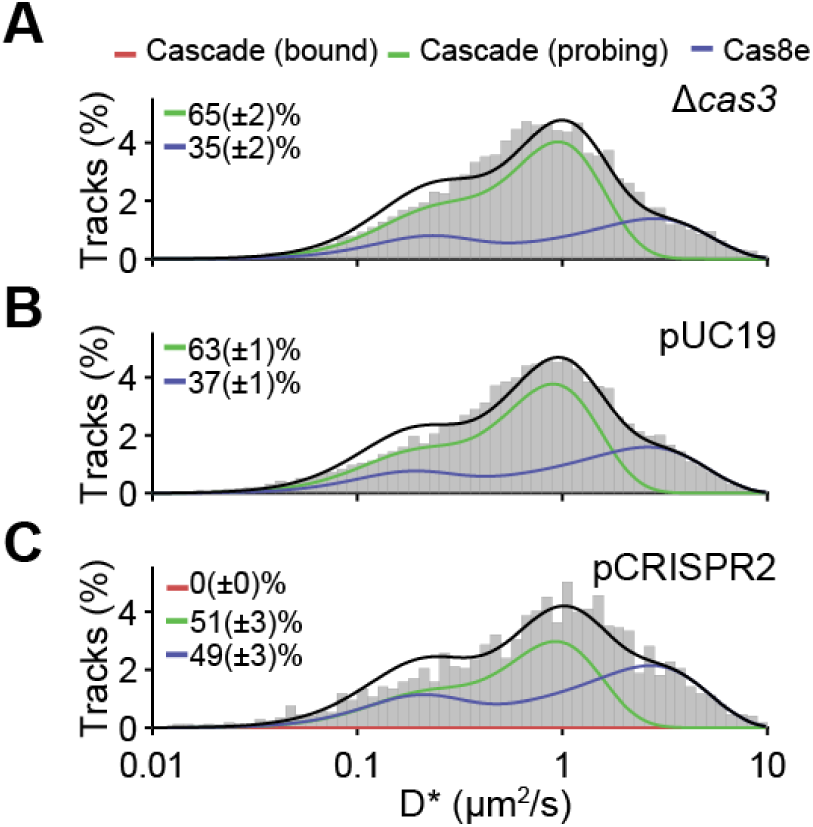
*D\** Histograms other conditions. *D*\* distributions for **(A)** Δ*cas3* strain, (**B**) Δ*cas3* strain + pUC19, the empty variant of pTarget-RNAP and pCRISPR1-RNAP and (**C**) Δ*cas3* strain + pCRISPR2. The amount of available Cascade complexes in the interference assay for strain pCRISPR2 targeting (Figure 5H) were extracted from the relative amount of Cascade complexes in this strain (51%) divided by the number of complexes in the WT strain (60%).

**Figure S4.**
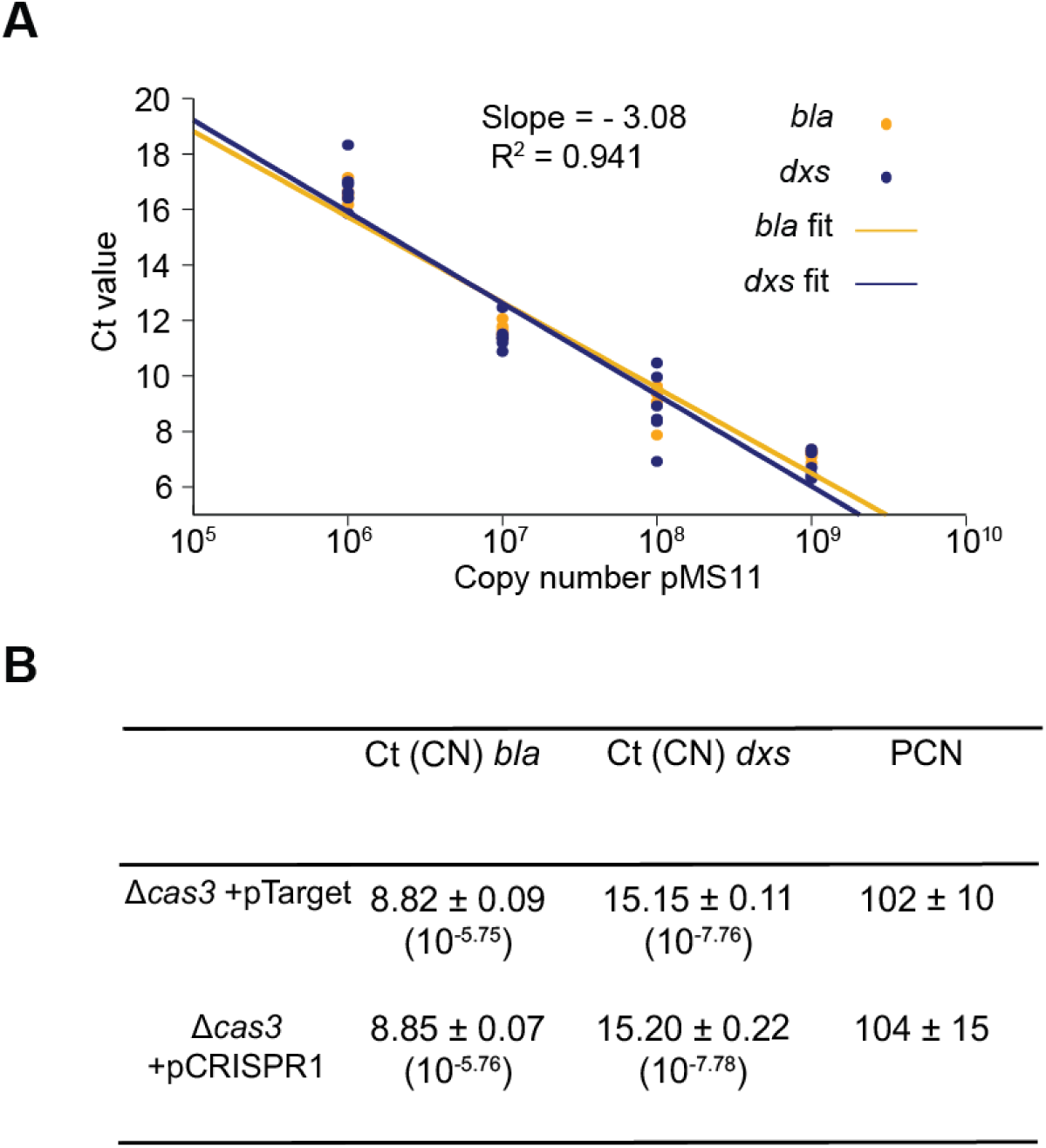
Plasmid copy number determination. **(A)** Calibration curve of *dxs* and *bla* primer amplification with dilution series of pMS11 (plasmid containing both *dxs* and *bla* gene). The regression of six technical replicates was used to make the calibration curve for both primer sets (regression parameters of *bla* and *dxs* gene in orange and purple respectively). (**B**) The Ct values of *bla* and *dxs* gene amplifications were calculated from biological triplicates. These Ct values were converted to absolute copy numbers (CN) by using the regression values from the calibration curve. The plasmid copy number per chromosome (PCN/chromosome) was calculated by dividing the copy number of the *bla* gene by the copy number of the *dxs* gene. The plasmid copy number per cell was estimated by multiplying PCN/chromosome by the expected number of chromosomes per cell (4) based on a literature value (Wallden et al., 2016).

**Figure S5.**
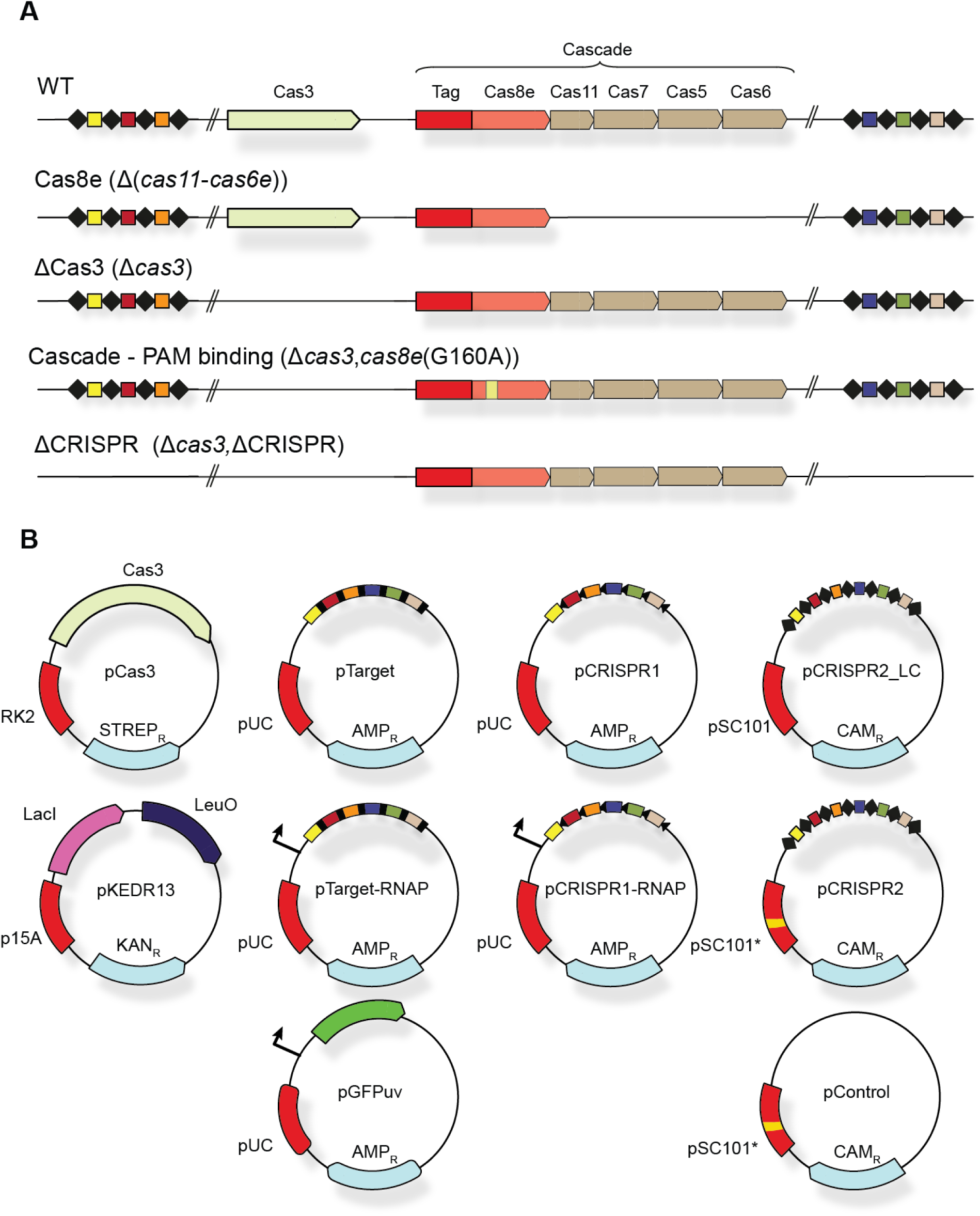
Strains and plasmids used. (**A**) Strains used in this study, strains were constructed with lambda recombination and verified by sequencing. Only part of each CRISPR array indicated (total 18 spacers). (**B**) Plasmids used in this study. Indicated are the ori (red), antibiotic resistance marker (light blue) and other components on the plasmid. Only part of the total 18 spacers are indicated for pTarget, pCRISPR1 and pCRISPR2. For sequences and descriptions see Table S3 and S4.

## Supplementary tables

**Table S1:**
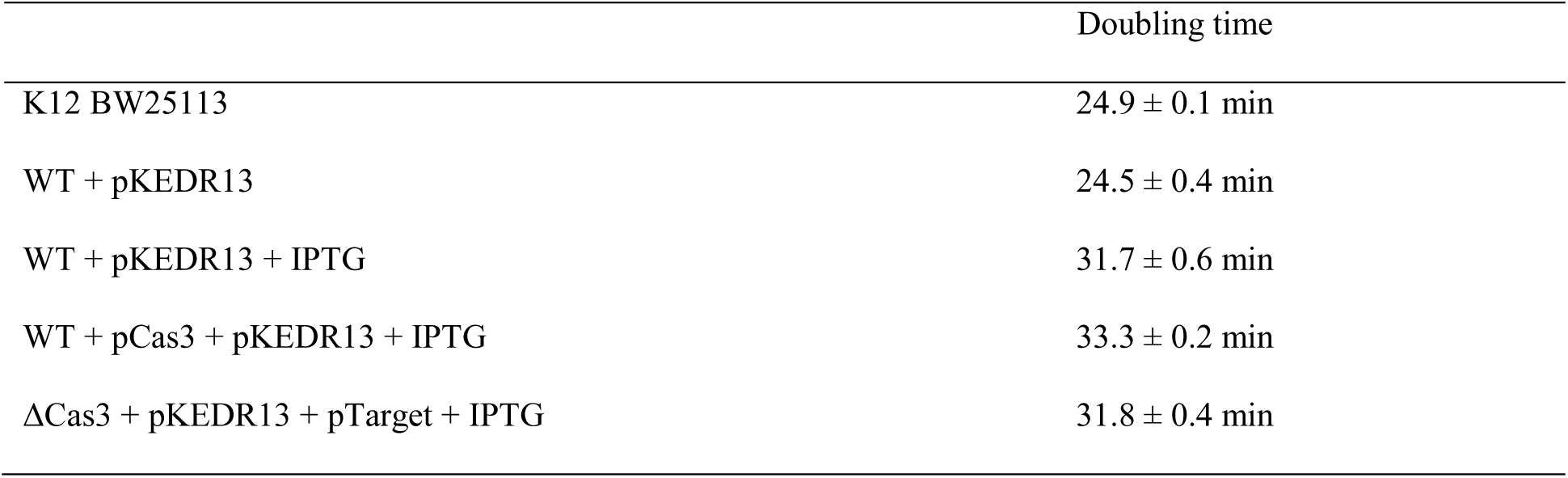
Growth rate of *E. coli* strains used in this study. Growth rates were determined in a plate reader where cells were inoculated in similar conditions as described in Methods. The instantaneous growth rate was determined at *t* = 2.5 hours, which represented the growth rate at the time of the microscope studies. Three independent cultures were measured to get the mean and standard error values.

|  | Doubling time |
| --- | --- |
| K12 BW25113 | 24.9 ± 0.1 min |
| WT + pKEDR13 | 24.5 ± 0.4 min |
| WT + pKEDR13 + IPTG | 31.7 ± 0.6 min |
| WT + pCas3 + pKEDR13 + IPTG | 33.3 ± 0.2 min |
| ΔCas3 + pKEDR13 + pTarget + IPTG | 31.8 ± 0.4 min |

**Table S2.**
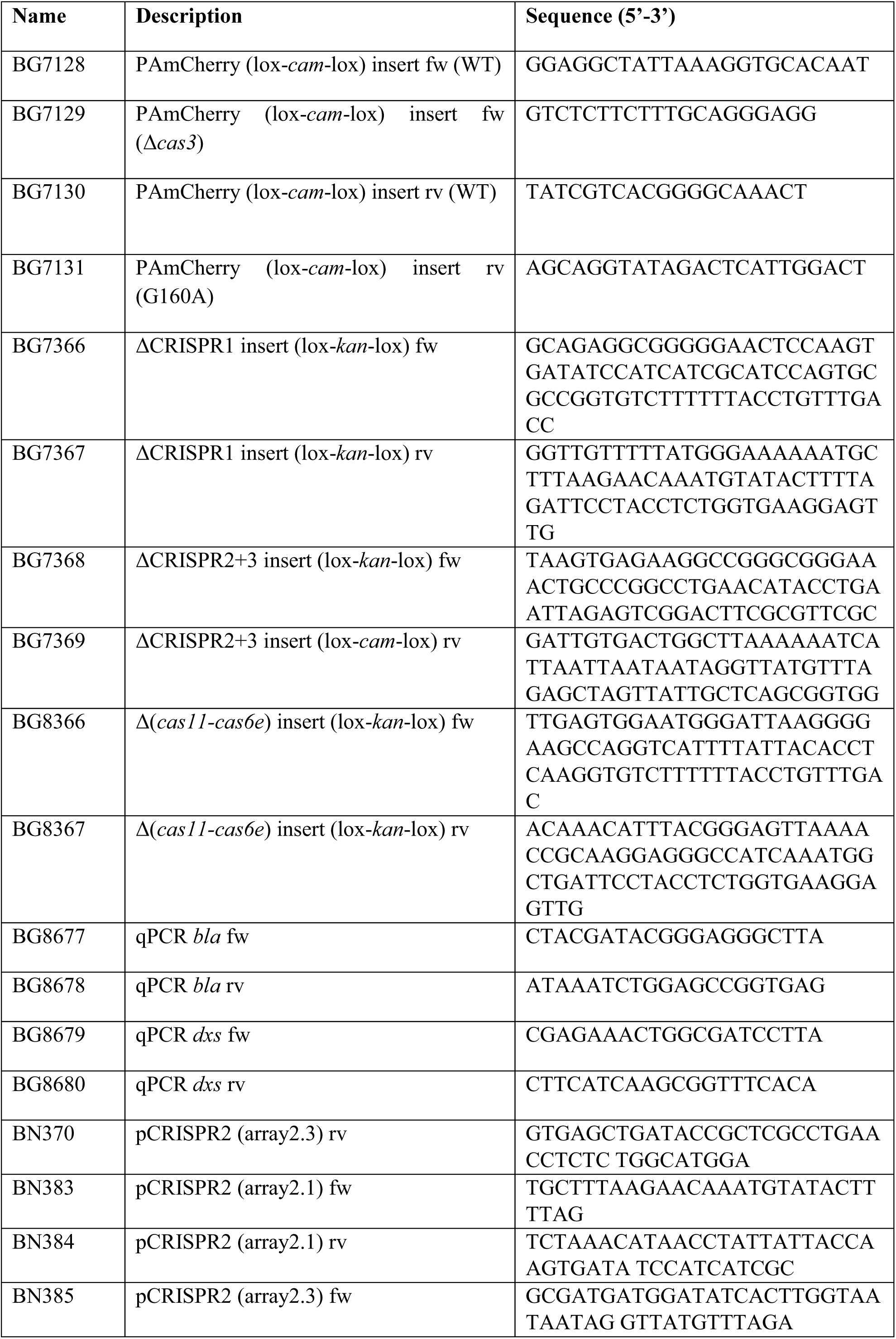
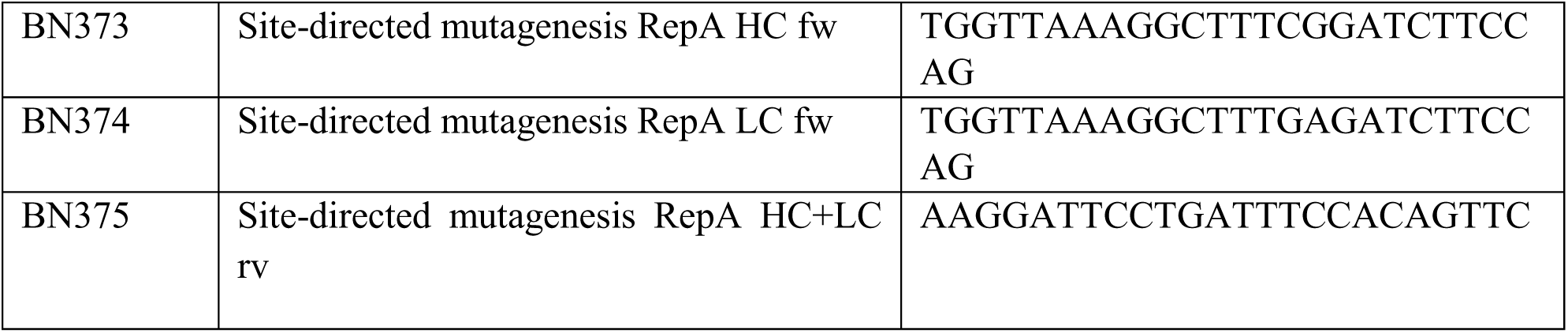
Primers used in this study.

**Table S3.**
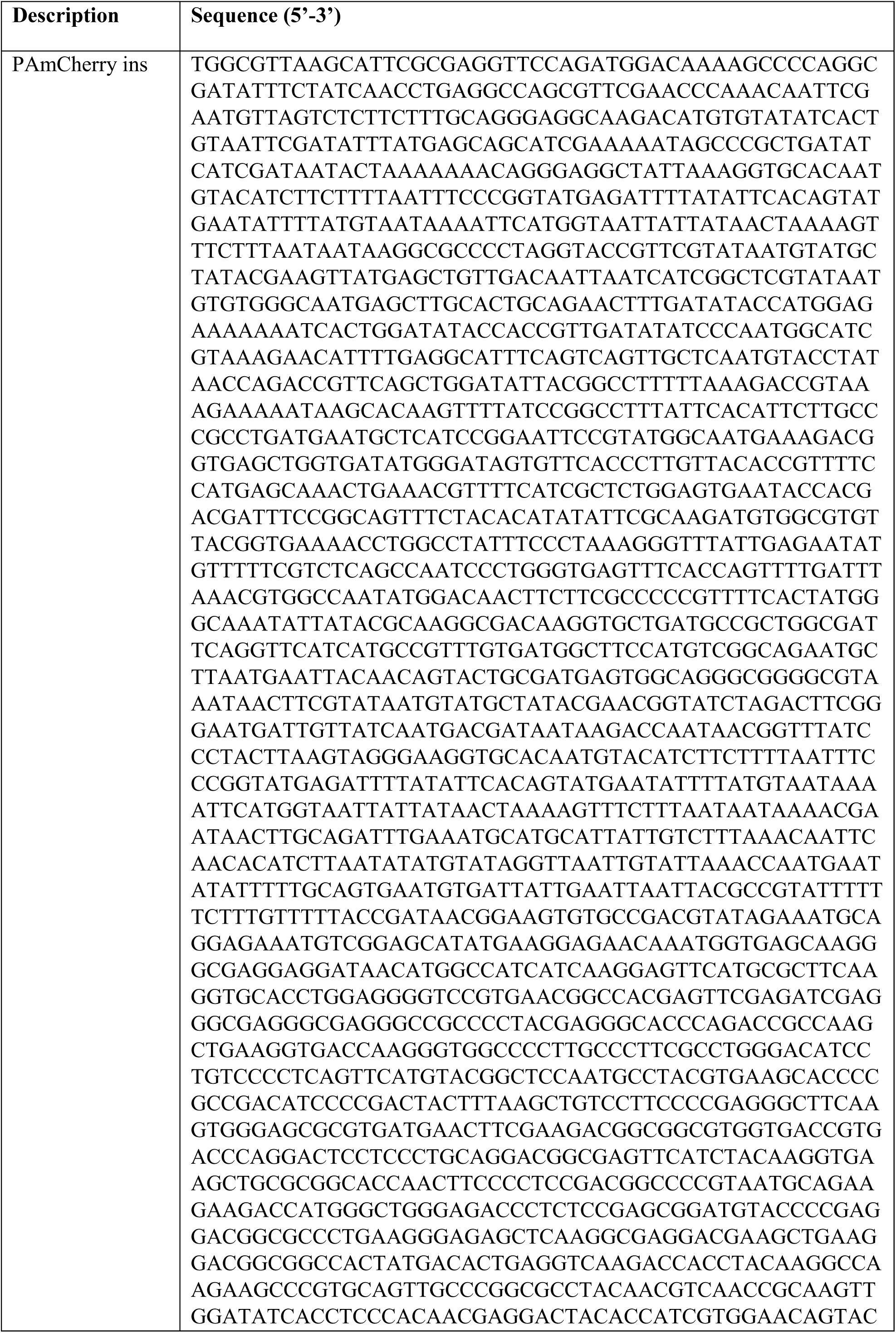
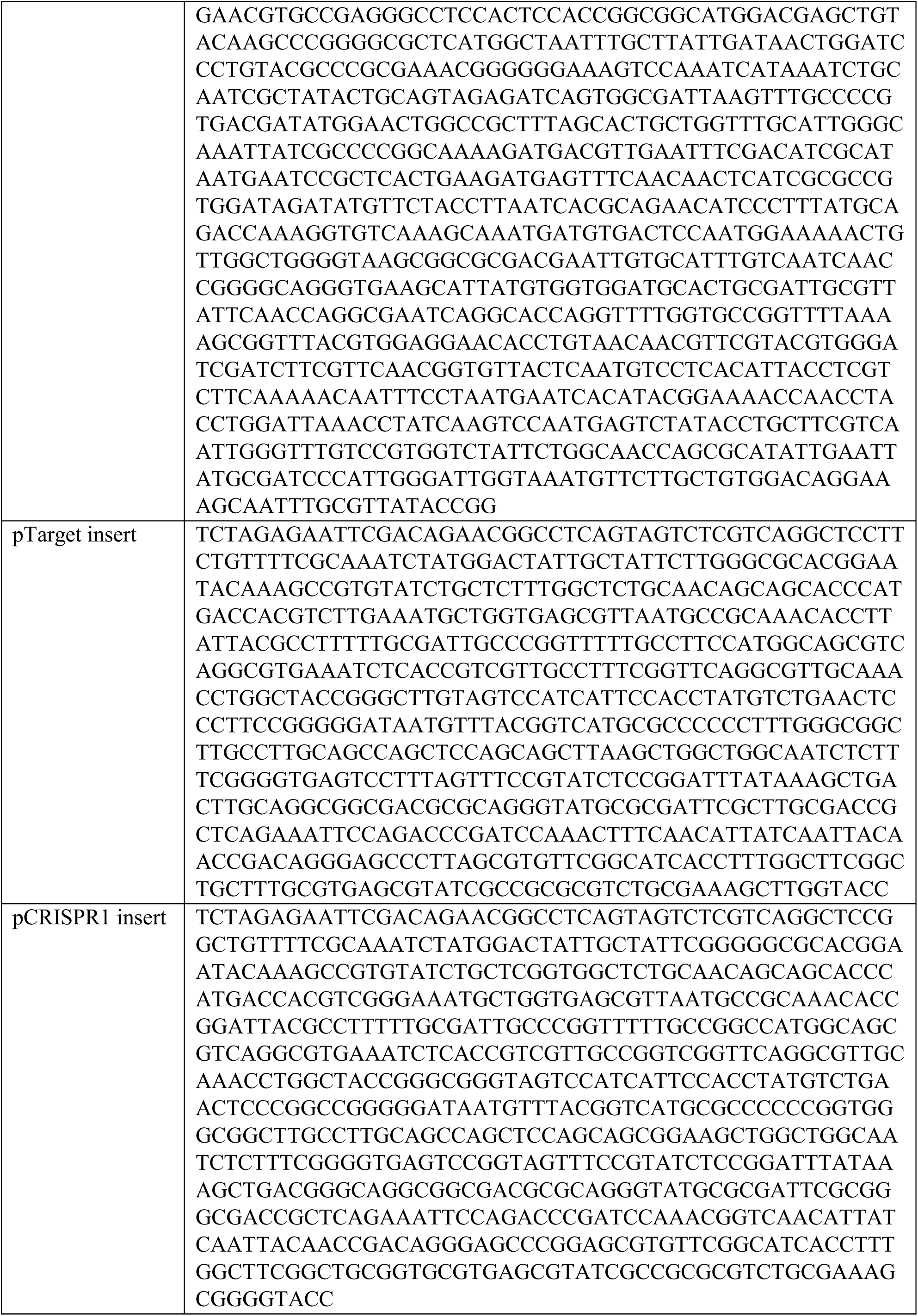
Synthetic DNA inserts used in this study.

**Table S4.** Plasmids used in this study.

| Name in study | Name in storage | Description | Source |
| --- | --- | --- | --- |
| pKEDR13 | pKEDR13 | Expression plasmid LeuO | (Westra et al., 2010) |
| pGFPuv | pGFPuv | Expression plasmid GFPuv | Clontech |
| pMS011 | pMS011 | Plasmid containing *bla* and *dxs* gene (qPCR) | (Caforio et al., 2018) |
| pSC020 | pSC020 | Plasmid containing Cre and lambda recombinase | S. Creutzberg (unpublished) |
| pTarget | pTU256 | Target plasmid containing all 18 potential protospacers for flanked by 5'-CTT-3' | This study |
| pTarget-RNAP | pTU150 | Target plasmid containing all 18 potential protospacers for flanked by 5'-CTT-3' and *plac* upstream | This study |
| pCRISPR1 | pTU258 | Target plasmid containing all 18 potential protospacers for flanked by 5'-CGG-3' | This study |
| pCRISPR1-RNAP | pTU152 | Target plasmid containing all 18 potential protospacers for flanked by 5'-CGG-3' and *plac* upstream | This study |
| pCRISPR2_LC | pTU158 | Plasmid containing all 18 potential protospacers for flanked by repeat sequences; low copy backbone variant of pSC101 | This study |
| pCRISPR2 | pTU160 | Plasmid containing all 18 potential protospacers for flanked by repeat sequences; high copy backbone variant of pSC101 | This study |
| pControl | pTU254 | High copy backbone variant of pSC101 | This study |
| pCas3 | pTU255 | Expression plasmid Cas3 | This study |

## Glossary

| Full name | Symbol | Description |
| --- | --- | --- |
| Apparent diffusion coefficient | $D^*$ | Apparent due to confinement |
| Bound state | $S1$ | |
| Dissociation constant | $K_D$ | Constant which is a measure for the binding affinity of two objects with each other |
| DNA segments | $sm_{DNA}$ | Amount of segments defined as containing DNA by DAPI staining |
| Enrichment Factor | $EF$ | The number of localizations in DNA-containing segments divided by the number of localizations in DNA-free segments |
| Fraction DNA bound | $f_{onDNA}$ | Fraction of the time DNA binding proteins spend on DNA is calculated from the off- and on-rate (Figure 1). |
| Frametime | $t_f$ | Positions of simulated/measured particles are recorded for each frametime |
| Free diffusion coefficient | $D_{free}^*$ | Diffusion coefficient in the absence of DNA binding. Apparent due to confinement. |
| Integrated time | $t_{int}$ | Overall timescale: can be one or multiple frametimes |
| Localization error | $\sigma$ | Average error in determination of particle position |
| Mobile state | $S2$ | |
| off-rate | $k_{off}$ | Rate DNA bound protein is released from DNA. Inverse of residence time |
| pseudo-first order on-rate | $k_{on}^*$ | Rate mobile protein is binding to DNA. As the amount of potential DNA probing sites is very large, on-rate is independent of DNA concentration (pseudo-first order) |
| pTarget establishment | $p_{establishment}$ | Measure for interference level calculated from the transformation ratio |
|  |  | of pTarget and pGFPuv (Eq. 1) |
| Time step | $t_{step}$ | Displacements of simulated particles are calculated for each time step |
| Total number of segments | $sm_{tot}$ | Total number of segments in the cell |

**Movie S1 Cascade diffusion through the cell.** Real-time data from WT strain where on left cumulative overlay of tracks (each track differently coloured) are plotted on top of a brightfield image showing the outline of the cells and on the right the fluorescent signal of the single molecules are depicted. Scale bar and time indicated at the bottom (total duration 50 s). The movie shows only a small part of a normal FOV.

## References

Antonik, M., Felekyan, S., Gaiduk, A., and Seidel, C.A.M. (2006). Separating structural heterogeneities from stochastic variations in fluorescence resonance energy transfer distributions via photon distribution analysis. J. Phys. Chem. B 110, 6970–6978.

Beloglazova, N., Kuznedelov, K., Flick, R., Datsenko, K. a., Brown, G., Popovic, a., Lemak, S., Semenova, E., Severinov, K., and Yakunin, a. F. (2015). CRISPR RNA binding and DNA target recognition by purified Cascade complexes from Escherichia coli. Nucleic Acids Res. 43, 530–543.

Blosser, T.R., Loeff, L., Westra, E.R., Vlot, M., Künne, T., Sobota, M., Dekker, C., Brouns, S.J.J., and Joo, C. (2015). Two distinct DNA binding modes guide dual roles of a CRISPR-Cas protein complex. Mol. Cell 58, 60–70.

Bondy-Denomy, J., Garcia, B., Strum, S., Du, M., Rollins, M.F., Hidalgo-Reyes, Y., Wiedenheft, B., Maxwell, K.L., and Davidson, A.R. (2015). Multiple mechanisms for CRISPR-Cas inhibition by anti-CRISPR proteins. Nature 526, 136–139.

Brouns, S.J.J., Jore, M.M., Lundgren, M., Westra, E.R., Slijkhuis, R.J.H., Snijders, A.P.L., Dickman, M.J., Makarova, K.S., Koonin, E. V, and van der Oost, J. (2008). Small CRISPR RNAs guide antiviral defense in prokaryotes. Science 321, 960–964.

Brown, M.W., Dillard, K.E., Xiao, Y., Dolan, A., Hernandez, E., Dahlhauser, S., Kim, Y., Myler, L.R., Anslyn, E., Ke, A., et al. (2018). Assembly and translocation of a CRISPR-Cas primed acquisition complex. Cell 175, 1–13.

Caforio, A., Siliakus, M.F., Exterkate, M., Jain, S., Jumde, V.R., Andringa, R.L.H., Kengen, S.W.M., Minnaard, A.J., Driessen, A.J.M., and van der Oost, J. (2018). Converting Escherichia coli into an archaebacterium with a hybrid heterochiral membrane. Proc. Natl. Acad. Sci. U. S. A. 115, 3704–3709.

Chandradoss, S.D., Haagsma, A.C., Lee, Y.K., Hwang, J.-H., Nam, J.-M., and Joo, C. (2014). Surface Passivation for Single-molecule Protein Studies. J. Vis. Exp. 86, 4–11.

Chen, Y.J., Wu, D., Gelbart, W., Knobler, C.M., Phillips, R., and Kegel, W.K. (2018). Two-Stage Dynamics of in Vivo Bacteriophage Genome Ejection. Phys. Rev. X 8.

Datsenko, K.A., and Wanner, B.L. (2000). One-step inactivation of chromosomal genes in Escherichia coli K-12 using PCR products. Proc. Natl. Acad. Sci. U. S. A. 97, 6640–6645.

Datsenko, K.A., Pougach, K., Tikhonov, A., Wanner, B.L., Severinov, K., and Semenova, E. (2012). Molecular memory of prior infections activates the CRISPR/Cas adaptive bacterial immunity system. Nat. Commun.

Davison, J. (2015). Pre-early functions of bacteriophage T5 and its relatives. Bacteriophage 5, e1086500.

Deveau, H., Barrangou, R., Garneau, J.E., Labonte, J., Fremaux, C., Boyaval, P., Romero, D.A., Horvath, P., and Moineau, S. (2008). Phage Response to CRISPR-Encoded Resistance in Streptococcus thermophilus. J. Bacteriol. 190, 1390–1400.

Durisic, N., Laparra-Cuervo, L., Sandoval-Álvarez, Á., Borbely, J.S., and Lakadamyali, M. (2014). Single-molecule evaluation of fluorescent protein photoactivation efficiency using an in vivo nanotemplate. Nat. Methods 11, 156–162.

Edelstein, A., Amodaj, N., Hoover, K., Vale, R., and Stuurman, N. (2010). Computer Control of Microscopes Using µManager. In Current Protocols in Molecular Biology, (Hoboken, NJ, USA: John Wiley & Sons, Inc.), p. Unit14.20.

English, B.P., Hauryliuk, V., Sanamrad, A., Tankov, S., Dekker, N.H., and Elf, J. (2011). Single-molecule investigations of the stringent response machinery in living bacterial cells. Proc. Natl. Acad. Sci. 108, 365–373.

Van Erp, P.B.G., Jackson, R.N., Carter, J., Golden, S.M., Bailey, S., and Wiedenheft, B. (2015). Mechanism of CRISPR-RNA guided recognition of DNA targets in Escherichia coli. Nucleic Acids Res. 43, 8381–8391.

Farooq, S., and Hohlbein, J. (2015). Camera-based single-molecule FRET detection with improved time resolution. Phys. Chem. Chem. Phys. 17, 27862–27872.

Floc’h, K., Lacroix, F., Barbieri, L., Servant, P., Galland, R., Butler, C., Sibarita, J.-B., Bourgeois, D., and Timmins, J. (2018). Bacterial cell wall nanoimaging by autoblinking microscopy. Sci. Rep. 8, 14038.

Gleditzsch, D., Pausch, P., Müller-Esparza, H., Özcan, A., Guo, X., Bange, G., and Randau, L. (2018). PAM identification by CRISPR-Cas effector complexes: diversified mechanisms and structures. RNA Biol. 15476286.2018.1504546.

Globyte, V., Lee, S.H., Bae, T., Kim, J., and Joo, C. (2018). CRISPR/Cas9 searches for a protospacer adjacent motif by lateral diffusion. EMBO J. e99466.

Hayes, R.P., Xiao, Y., Ding, F., van Erp, P.B.G., Rajashankar, K., Bailey, S., Wiedenheft, B., and Ke, A. (2016). Structural basis for promiscuous PAM recognition in type I–E Cascade from E. coli. Nature 530, 499–503.

Ho, H.N., Van Oijen, A.M., and Ghodke, H. (2018). The transcription-repair coupling factor Mfd associates with RNA polymerase in the absence of exogenous damage. Nat. Commun. 9, 1570.

Hochstrasser, M.L., Taylor, D.W., Bhat, P., Guegler, C.K., Sternberg, S.H., Nogales, E., and Doudna, J. a. (2014). CasA mediates Cas3-catalyzed target degradation during CRISPR RNA-guided interference. Proc. Natl. Acad. Sci. 111, 6618–6623.

Holden, S.J., Uphoff, S., Hohlbein, J., Yadin, D., Le Reste, L., Britton, O.J., and Kapanidis, A.N. (2010). Defining the Limits of Single-Molecule FRET Resolution in TIRF Microscopy. Biophys. J. 99, 3102–3111.

Hoogendoorn, E., Crosby, K.C., Leyton-Puig, D., Breedijk, R.M.P., Jalink, K., Gadella, T.W.J., and Postma, M. (2015). The fidelity of stochastic single-molecule super-resolution reconstructions critically depends upon robust background estimation. Sci. Rep. 4, 3854.

Høyland-Kroghsbo, N.M., Muñoz, K.A., and Bassler, B.L. (2018). Temperature, by Controlling Growth Rate, Regulates CRISPR-Cas Activity in Pseudomonas aeruginosa. MBio 9.

Huang, B., Wu, H.K., Bhaya, D., Grossman, A., Granier, S., Kobilka, B.K., Zare, R.N., Huang, B., Wu, H.K., Bhaya, D., et al. (2007). Counting low-copy number proteins in a single cell. Science 315, 81–84.

Huang, F., Hartwich, T.M.P., Rivera-Molina, F.E., Lin, Y., Duim, W.C., Long, J.J., Uchil, P.D., Myers, J.R., Baird, M.A., Mothes, W., et al. (2013). Video-rate nanoscopy using sCMOS camera-specific single-molecule localization algorithms. Nat. Methods 10, 653–658.

Jackson, S.A., McKenzie, R.E., Fagerlund, R.D., Kieper, S.N., Fineran, P.C., and Brouns, S.J.J. (2017). CRISPR-Cas: Adapting to change. Science (80-.).

Jones, D.L., Leroy, P., Unoson, C., Fange, D., Ćurić, V., Lawson, M.J., and Elf, J. (2017). Kinetics of dCas9 target search in Escherichia coli. Science 357, 1420–1424.

Jore, M.M., Lundgren, M., van Duijn, E., Bultema, J.B., Westra, E.R., Waghmare, S.P., Wiedenheft, B., Pul, Ü., Wurm, R., Wagner, R., et al. (2011). Structural basis for CRISPR RNA-guided DNA recognition by Cascade. Nat. Struct. Mol. Biol. 18, 529–536.

Jung, C., Hawkins, J.A., Jones, S.K., Xiao, Y., Rybarski, J.R., Dillard, K.E., Hussmann, J., Saifuddin, F.A., Savran, C.A., Ellington, A.D., et al. (2017). Massively Parallel Biophysical Analysis of CRISPR-Cas Complexes on Next Generation Sequencing Chips. Cell 170, 35–47.e13.

Kalinin, S., Felekyan, S., Valeri, A., and Seidel, C.A.M. (2008). Characterizing Multiple Molecular States in Single-Molecule Multiparameter Fluorescence Detection by Probability Distribution Analysis. J. Phys. Chem. B 112, 8361–8374.

Knight, S.C., Xie, L., Deng, W., Guglielmi, B., Witkowsky, L.B., Bosanac, L., Zhang, E.T., El Beheiry, M., Masson, J.-B.J.-B.J.-B., Dahan, M., et al. (2015). Dynamics of CRISPR-Cas9 genome interrogation in living cells. Science 350, 823–826.

Kumar, M., Mommer, M.S., and Sourjik, V. (2010). Mobility of cytoplasmic, membrane, and DNA-binding proteins in Escherichia coli. Biophys. J. 98, 552–559.

Lee, S.-H., Shin, J.Y., Lee, A., and Bustamante, C. (2012). Counting single photoactivatable fluorescent molecules by photoactivated localization microscopy (PALM). Proc. Natl. Acad. Sci. 109, 17436–17441.

Leenay, R.T., Maksimchuk, K.R., Slotkowski, R.A., Agrawal, R.N., Gomaa, A.A., Briner, A.E., Barrangou, R., and Beisel, C.L. (2016). Identifying and Visualizing Functional PAM Diversity across CRISPR-Cas Systems. Mol. Cell 62, 137–147.

Ma, J., and Wang, M.D. (2016). DNA supercoiling during transcription. Biophys. Rev. 8, 75–87.

Majsec, K., Bolt, E.L., and Ivančić-Baće, I. (2016). Cas3 is a limiting factor for CRISPR-Cas immunity in Escherichia coli cells lacking H-NS. BMC Microbiol. 16, 28.

Manley, S., Gillette, J.M., Patterson, G.H., Shroff, H., Hess, H.F., Betzig, E., and Lippincott-Schwartz, J. (2008). High-density mapping of single-molecule trajectories with photoactivated localization microscopy. Nat. Methods 5, 155–157.

Marraffini, L.A. (2015). CRISPR-Cas immunity in prokaryotes. Nature 526, 55–61.

Martens, K.J.A., Beljouw, S. van Els, S. van der Baas, S., Vink, J.N.A., Brouns, S.J.J., Baarlen, P. van Kleerebezem, M., and Hohlbein, J. (2018). An open microscopy framework suited for tracking dCas9 in live bacteria. BioRxiv 437137.

Martynov, A., Severinov, K., and Ispolatov, I. (2017). Optimal number of spacers in CRISPR arrays. PLoS Comput. Biol. 13.

McGuigan, J.A.S., Kay, J.W., and Elder, H.Y. (2006). Critical review of the methods used to measure the apparent dissociation constant and ligand purity in Ca2+ and Mg2+ buffer solutions. Prog. Biophys. Mol. Biol. 92, 333–370.

Michalet, X. (2010). Mean square displacement analysis of single-particle trajectories with localization error: Brownian motion in an isotropic medium. Phys. Rev. E - Stat. Nonlinear, Soft Matter Phys. 82, 041914.

Mika, J.T., and Poolman, B. (2011). Macromolecule diffusion and confinement in prokaryotic cells. Curr. Opin. Biotechnol. 22, 117–126.

Mika, J.T., Van Den Bogaart, G., Veenhoff, L., Krasnikov, V., and Poolman, B. (2010). Molecular sieving properties of the cytoplasm of Escherichia coli and consequences of osmotic stress. Mol. Microbiol. 77, 200–207.

Modell, J.W., Jiang, W., and Marraffini, L.A. (2017). CRISPR-Cas systems exploit viral DNA injection to establish and maintain adaptive immunity. Nature 544, 101–104.

Mojica, F.J.M., Díez-Villaseñor, C., García-Martínez, J., and Almendros, C. (2009). Short motif sequences determine the targets of the prokaryotic CRISPR defence system. Microbiology 155, 733–740.

Mondal, J., Bratton, B.P., Li, Y., Yethiraj, A., and Weisshaar, J.C. (2011). Entropy-based mechanism of ribosome-nucleoid segregation in E. coli Cells. Biophys. J. 100, 2605–2613.

Nenninger, A., Mastroianni, G., and Mullineaux, C.W. (2010). Size dependence of protein diffusion in the cytoplasm of Escherichia coli. J. Bacteriol. 192, 4535–4540.

Olsson, J.A., Berg, O.G., Dasgupta, S., and Nordström, K. (2003a). Eclipse period during replication of plasmid R1: contributions from structural events and from the copy-number control system. Mol. Microbiol. 50, 291–301.

Olsson, J.A., Berg, O.G., Dasgupta, S., and Nordström, K. (2003b). Eclipse period during replication of plasmid R1: contributions from structural events and from the copy-number control system. Mol. Microbiol. 50, 291–301.

Paintdakhi, A., Parry, B., Campos, M., Irnov, I., Elf, J., Surovtsev, I., and Jacobs-Wagner, C. (2016). Oufti: An integrated software package for high-accuracy, highthroughput quantitative microscopy analysis. Mol. Microbiol. 99, 767–777.

Palo, K., Mets, Ü., Loorits, V., and Kask, P. (2006). Calculation of photon-count number distributions via master equations. Biophys. J. 90, 2179–2191.

Pawluk, A., Bondy-Denomy, J., Cheung, V.H.W., Maxwell, K.L., and Davidson, A.R. (2014). A new group of phage anti-CRISPR genes inhibits the type I-E CRISPR-Cas system of pseudomonas aeruginosa. MBio 5.

Peterson, J., and Phillips, G.J. (2008). New pSC101-derivative cloning vectors with elevated copy numbers. Plasmid 59, 193–201.

Qian, H., Sheetz, M.P., and Elson, E.L. (1991). Single particle tracking. Analysis of diffusion and flow in two-dimensional systems. Biophys. J. 60, 910–921.

Redding, S., Sternberg, S.H.H., Marshall, M., Gibb, B., Bhat, P., Guegler, C.K.K., Wiedenheft, B., Doudna, J.A., and Greene, E.C.C. (2015). Surveillance and Processing of Foreign DNA by the Escherichia coli CRISPR-Cas System. Cell 163, 1–12.

Reyes-Lamothe, R., Tran, T., Meas, D., Lee, L., Li, A.M., Sherratt, D.J., and Tolmasky, M.E. (2014). High-copy bacterial plasmids diffuse in the nucleoid-free space, replicate stochastically and are randomly partitioned at cell division. Nucleic Acids Res. 42, 1042–1051.

Sanamrad, A., Persson, F., Lundius, E.G., Fange, D., Gynna, A.H., and Elf, J. (2014). Single-particle tracking reveals that free ribosomal subunits are not excluded from the Escherichia coli nucleoid. Proc. Natl. Acad. Sci. 111, 11413–11418.

Sashital, D.G., Wiedenheft, B., and Doudna, J.A. (2012). Mechanism of Foreign DNA Selection in a Bacterial Adaptive Immune System. Mol. Cell 46, 606–615.

Severinov, K., Ispolatov, I., and Semenova, E. (2016). The Influence of Copy-Number of Targeted Extrachromosomal Genetic Elements on the Outcome of CRISPR-Cas Defense. Front. Mol. Biosci. 3.

Shao, Q., Hawkins, A., and Zeng, L. (2015). Phage DNA Dynamics in Cells with Different Fates. Biophys. J. 108, 2048–2060.

De Smet, J., Hendrix, H., Blasdel, B.G., Danis-Wlodarczyk, K., and Lavigne, R. (2017). Pseudomonas predators: Understanding and exploiting phage-host interactions. Nat. Rev. Microbiol.

Staals, R.H.J., Jackson, S.A., Biswas, A., Brouns, S.J.J., Brown, C.M., and Fineran, P.C. (2016). Interference-driven spacer acquisition is dominant over naive and primed adaptation in a native CRISPR-Cas system. Nat. Commun. 7.

Sternberg, S.H., Redding, S., Jinek, M., Greene, E.C., and Doudna, J.A. (2014). DNA interrogation by the CRISPR RNA-guided endonuclease Cas9. Nature 507, 62–67.

Stracy, M., Lesterlin, C., Garza de Leon, F., Uphoff, S., Zawadzki, P., and Kapanidis, A.N. (2015). Live-cell superresolution microscopy reveals the organization of RNA polymerase in the bacterial nucleoid. Proc. Natl. Acad. Sci. 112, E4390–E4399.

Subach, F. V., Patterson, G.H., Manley, S., Gillette, J.M., Lippincott-Schwartz, J., and Verkhusha, V. V. (2009). Photoactivatable mCherry for high-resolution two-color fluorescence microscopy. Nat. Methods 6, 153–159.

Szczelkun, M.D., Tikhomirova, M.S., Sinkunas, T., Gasiunas, G., Karvelis, T., Pschera, P., Siksnys, V., and Seidel, R. (2014). Direct observation of R-loop formation by single RNA-guided Cas9 and Cascade effector complexes. Proc. Natl. Acad. Sci. 111, 9798–9803.

Uphoff, S., Reyes-Lamothe, R., Garza de Leon, F., Sherratt, D.J., and Kapanidis, A.N. (2013). Single-molecule DNA repair in live bacteria. Proc. Natl. Acad. Sci. U. S. A. 110, 8063–8068.

Vigouroux, A., Oldewurtel, E., Cui, L., Bikard, D., and van Teeffelen, S. (2018). Tuning dCas9’s ability to block transcription enables robust, noiseless knockdown of bacterial genes. Mol. Syst. Biol. 14, e7899.

Vliet, L. Van, Sudar, D., and Young, I. (1998). Digital fluorescence imaging using cooled charge-coupled device array cameras. Cell Biol. III, 109–120.

Vrljic, M., Nishimura, S.Y., Brasselet, S., Moerner, W.E., and McConnell, H.M. (2002). Translational diffusion of individual class II MHC membrane proteins in cells. Biophys. J. 83, 2681–2692.

Wallden, M., Fange, D., Lundius, E.G., Baltekin, Ö., and Elf, J. (2016). The Synchronization of Replication and Division Cycles in Individual E. coli Cells. Cell 166, 729–739.

Westra, E.R., Pul, Ü., Heidrich, N., Jore, M.M., Lundgren, M., Stratmann, T., Wurm, R., Raine, A., Mescher, M., Van Heereveld, L., et al. (2010). H-NS-mediated repression of CRISPR-based immunity in Escherichia coli K12 can be relieved by the transcription activator LeuO. Mol. Microbiol. 77, 1380–1393.

Westra, E.R., van Erp, P.B.G., Künne, T., Wong, S.P., Staals, R.H.J., Seegers, C.L.C., Bollen, S., Jore, M.M., Semenova, E., Severinov, K., et al. (2012). CRISPR Immunity Relies on the Consecutive Binding and Degradation of Negatively Supercoiled Invader DNA by Cascade and Cas3. Mol. Cell 46, 595–605.

Xiao, Y., Luo, M., Hayes, R.P., Kim, J., Ng, S., Ding, F., Liao, M., and Ke, A. (2017). Structure Basis for Directional R-loop Formation and Substrate Handover Mechanisms in Type I CRISPR-Cas System. Cell 170, 48–60.

Xiao, Y., Luo, M., Dolan, A.E., Liao, M., and Ke, A. (2018). Structure basis for RNA-guided DNA degradation by Cascade and Cas3. Science 361, eaat0839.

Xue, C., Whitis, N.R., and Sashital, D.G. (2016). Conformational Control of Cascade Interference and Priming Activities in CRISPR Immunity. Mol. Cell 64, 826–834.

Xue, C., Zhu, Y., Zhang, X., Shin, Y.K., and Sashital, D.G. (2017). Real-Time Observation of Target Search by the CRISPR Surveillance Complex Cascade. Cell Rep. 21, 3717–3727.

Zawadzki, P., Stracy, M., Ginda, K., Zawadzka, K., Lesterlin, C., Kapanidis, A.N., and Sherratt, D.J. (2015). The Localization and Action of Topoisomerase IV in Escherichia coli Chromosome Segregation Is Coordinated by the SMC Complex, MukBEF. Cell Rep. 13, 2587–2596.

